# Termite-Like Phylogenetic Conservation of Gut Microbiota in Passalid Beetles Suggests Evolutionary Convergence in Decaying Wood Degradation

**DOI:** 10.64898/2026.10.28.762520

**Authors:** Cristian Fernando Beza-Beza, Larry Antonio Jiménez-Ferbans, Andrés Redondo-Soto, Michael Vincent Welsh, Aram Mikaelyan

## Abstract

Beetles of the family Passalidae are an important group of subsocial wood-decomposing (saproxylophagous) insects, spending their entire life cycle living inside and feeding on rotten logs. Decaying wood is an abundant resource, but nutritionally poor and challenging to digest. As in other saproxylophagous insects, digestion for passalids is only possible through symbiotic relationships with their gut microbiota, highlighting the critical roles of these symbionts in the evolution of wood-decomposing insects. Despite its importance, this critical symbiotic relationship remains poorly understood, hindering our ability to identify the factors shaping this association and assessing its role in the evolution of Passalidae. Using high-throughput sequencing of the microbial 16S rRNA we comparatively analyzed the composition and abundance of symbiont lineages in the anterior hindguts of passalid beetles from two independent mountain systems in the Colombian Caribbean region. We tested the influence of geographical co-occurrence and phylogenetic relatedness on gut microbial composition and structure. We posit that if shared locality was the primary driver of gut microbiome assembly, then sympatric species should harbor more similar microbiomes regardless of their phylogenetic relationships. Alternatively, if host evolutionary history is the dominant factor, congeneric species should exhibit more similar microbiomes regardless of geographic co-occurrence. Our results support that phylogenetic relatedness explained substantially more variation in both bacterial and archaeal communities than geographic locality, indicating that gut microbiome composition is primarily structured by host evolutionary history. Furthermore, bacterial and archaeal communities exhibited evidence of phylosymbiosis, although at different phylogenetic scales: bacterial communities more closely reflected relationships among genus level clades, whereas archaeal communities retained congruence mainly at deeper host lineages. We also uncovered extensive bacterial taxonomic novelty, with nearly half of bacterial reads remaining unclassified at the genus level, highlighting the large undescribed microbial diversity associated with Passalidae. These findings lead to establishing Passalidae as a promising comparative model for investigating the evolutionary processes governing gut microbiome assembly and the evolution of lignocellulose-feeding symbioses.

## Introduction

Wood is an exceptionally demanding diet: it is nitrogen-poor, physically defended, and requires specialized physiological and microbial machinery to access (Beza-Beza et al., 2024). Only a few insect lineages have independently evolved the ability to digest lignocellulose internally, these independent origins provide natural replicates for testing whether saproxylophagy predictably produces microbiomes that track host evolutionary history, or whether similar systems of symbiotic digestion can be sustained by microbial communities that turn over across hosts and environments. Termites, wood-feeding cockroaches, and passalid beetles represent independent solutions to this challenge, with gut microbiomes that serve broadly similar functions but differ markedly in structure and in how they are assembled and maintained (Beza-Beza et al., 2024). Termites exhibit evidence for long-term host–microbiome fidelity known in insects, including multi-lineage symbiont clades whose phylogenies parallel host diversification over tens of millions of years (Arora et al., 2023; Bourguignon et al., 2018). By contrast, wood-feeding cockroaches show strong host-species clustering but only weak or inconsistent phylosymbiosis once deeper evolutionary relationships are considered, with ecological filtering and gut physicochemistry overriding host phylogeny (Bauer et al., 2015; Dietrich et al., 2014; Tinker & Ottesen, 2020).

Passalid beetles (Coleoptera: Passalidae) represent another independent origin of gut-based decaying-wood feeding. With close to a 1,000 described species (Boucher, 2006), they are one of the principal wood decomposers in the tropics due to their entire wood-dwelling lifestyle and large size, corresponding to higher food intake (Castillo & Reyes-Castillo, 2009). For example, some species have been estimated to process up to 4.5 times their own body weight in wood per day (Castillo & Morón, 1992); and yet, their microbiome evolution remains largely unexplored. Existing work, mainly limited to *Odontotaenius disjunctus*, reveals a compartmentalized gut with predictable physicochemical structure and a microbe-rich frass provisioning system that larvae rely on for nutrition and microbial inoculation (Ceja-Navarro et al., 2014, 2019; Schwarz et al., 2023). These features, together with subsociality and repeated use of the same galleries, create conditions expected to promote partner fidelity and lineage-specific microbial associations. At the same time, several functionally important passalid gut taxa appear to be habitat-defined rather than host-defined, including fiber-attached bacterial consortia and recurrent defensive actinomycetes found in frass and pupal chambers across broad geographic ranges (Schwarz et al., 2023). Together, these features position Passalidae as the only available system outside termites, in which competing hypotheses about host-driven versus environmentally assembled microbiomes can be directly tested. Despite this, virtually nothing is known about gut microbial variation across passalid species, genera, or geographic space. Whether gut community similarity tracks host phylogeny across Passalidae (phylosymbiosis) remains unresolved. It is also unclear whether distantly related passalid species share bacterial or archaeal clades and whether shared geographic distribution overwhelms or reinforces patterns associated with host filtering. This gap is particularly pronounced for Archaea, which have never been profiled across multiple passalid hosts, despite evidence from other insects that methanogenic archaeal distributions are strongly constrained by gut physicochemistry, including H₂ availability, pH, and O₂ tension (Egert et al., 2003; Lemke et al., 2003).

Across decomposer insects, contrasting patterns further underscore the need for comparative datasets. In bark beetles and scarabs, geographic turnover is common but often accompanied by functionally stable microbial guilds (Han et al., 2024; Hernández-García et al., 2018). In omnivorous cockroaches, microbiomes strongly cluster by species yet exhibit only weak phylosymbiosis (Tinker & Ottesen, 2020). Comparative work illustrates a broader principle: phylosymbiosis emerges when vertical transmission and mutualistic dependence dominate, and erodes when environmental acquisition, ecological filtering, or permissive partner choice overshadow host lineage effects (Hayward et al., 2021; Lim & Bordenstein, 2020). Termites exemplify the first pattern; wood-feeding cockroaches and cerambycids often exemplify the second.

Together, these observations highlight a major gap in our understanding of gut-microbiome evolution in insect wood-feeders: although passalids have a relatively well-resolved digestive physiology and plausible routes of symbiont transfer, we do not know whether their microbiome is evolutionarily conserved, ecologically assembled, or whether some combination of the two exists. No study has jointly examined multiple sympatric species, across multiple sites, using standardized methods to test phylosymbiosis, identify core taxa, or quantify bacterial versus archaeal structuring. As a result, we lack resolution on whether passalid beetles behave more like termites—showing host-tracking, lineage-specific microbial partners—or more like cockroaches, where gut physicochemistry and ecological filtering dominate microbiome composition and structure.

In this study, we test whether an independent origin of wood-feeding recapitulates the evolutionary patterns observed in termites or instead follows the more environmentally structured dynamics seen in cockroaches. By sampling multiple passalid species across two biogeographically distinct mountain systems, we integrate host phylogenetic reconstruction with bacterial and archaeal community profiling to evaluate how microbiome variation partitions across evolutionary and geographic scales. We specifically ask whether microbial communities track host lineage, whether this signal differs between bacteria and archaea, and whether geographic separation disrupts or reinforces host-associated structure. This framework allows us to distinguish between phylogenetic niche conservatism, ecological assembly, and mixed-mode dynamics in a lineage where these processes have never been evaluated comparatively.

## Methods

### Study area and main research design

Species of the family Passalidae exhibit a high degree of endemism and elevational specialization (Beza-Beza et al., 2021). Taking advantage of this characteristic, we have designed an experiment to test phylosymbioses in the group. We selected two localities in Colombia, separated by a dry valley, where we can find the same genera but different species of each. Specimens were collected from two locations in the Sierra Nevada de Santa Marta (altitudes 1662–1684 m, separated by 750 m) and four locations in the Serranía del Perijá (altitudes 1745–1792 m, separated by a maximum distance of 340 m) (Fig. 1). At each location, we examine rotten logs during free-ranging surveys. The specimens were identified using the taxonomic identification key in Jiménez-Ferbans et al. (2023), and comparison with material deposited at the Colección Entomológica del Centro de Coleciones Biológicas (Universidad del Magdalena). See Supplementary Table S1 for detailed specimen information.

**Figure 1.**
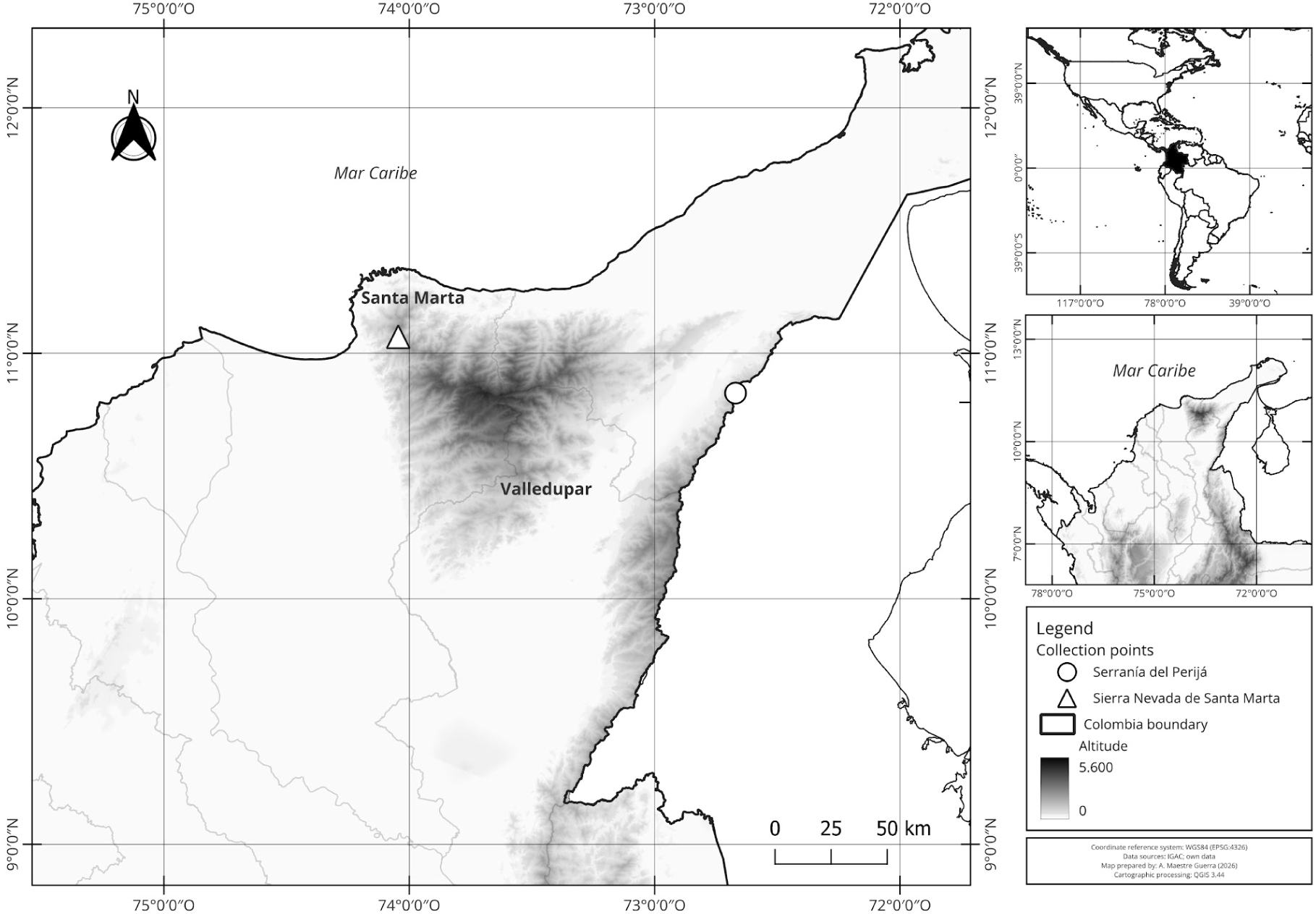
Map of area of study showing the topographic profile of the zone and the two collecting localities.

To test the effect of evolutionary history on the microbial composition at different levels of phylogenetic depth, we sampled members of two tribes and four genera of bess beetles. New World passalids are represented by the tribes Proculini and Passalini, each comprising approximately 200 described species. According to Beza-Beza et al. (2021), both tribes are reciprocally monophyletic and diverged between 112 and 135 Ma. Passalini is particularly diverse in South America, whereas Proculini reaches its highest diversity in Mesoamerica and is mainly distributed across montane systems associated with the Andes (Beza-Beza et al., 2021; Jiménez-Ferbans et al., 2018). In order to achieve an adequate phylogenetic and taxonomic representation, we selected two genera of Passalini (*Passalus* and *Rhodocanthopus*) and two of Proculini (*Popilius* and *Veturius*) (Table 1). Based on the estimates of Beza-Beza et al. (2021), the most recent common ancestor of *Passalus* and *Rhodocanthopus* lived approximately 60 Ma, whereas the lineage leading to *Popilius* and *Veturius* diverged around 50 Ma.

**Table 1.** Distribution and elevational range of Passalidae species sampled. The elevational ranges provided are local to each mountain system, and when the species is distributed more broadly, species level elevation ranges are given in parentheses (taken from Jiménez-Ferbans et al., 2023b).

| Species | Distribution | Elevational range (m) |
|---|---|---|
| S.N.S.M. |  |  |
| *Passalus* (*Passalus*) *coniferus* Eschscholtz | Argentina-northern Colombia. | 710–1880 (1550–1560) |
| *Passalus*. (*Pertinax*) *unimagdalenae* Jiménez-Ferbans, Reyes-Castillo & Amat-García | Endemic to S.N.S.M. | 710–2260 (1560–2309) |
| *Popilius* aff. *marginatus* | S.N.S.M. | 870–1900 |
| *Veturius* (*Publius*) *impressus* Hincks | Endemic to S.N.S.M. | 1270–2050 (1560) |
| **Serranía del Perijá** |  |  |
| *Rhodocanthopus maillei* (Percheron) | Northern Andes | 980–2280 (180–2400) |
| *Popilius gibbosus* (Burmeister) | Cordillera Oriental of the northern Andes | 1520–2470 (1350–3000) |
| *Veturius* (*Veturius*) *standfussi* Kuwert | Andes of Bolivia-Venezuela | 1280–1910 (800–2500) |
| *Passalus* (*Passalus*) *punctiger* Lepeletier & Serville | Argentina-Mexico | 20–2260 (30–1500) |

### DNA extraction and amplification of microbial communities

We transported live specimens from the field in plastic containers along with pieces of wood obtained from the logs in which the specimens were found, and kept them overnight at the Laboratorio de Biodiversidad, Sistemática y Biogeografía at Universidad del Magdalena, Santa Marta, Colombia. Following the waiting period, we performed dissections to extract the anterior hindgut (AHG), which houses the majority of the bacterial load in *O. disjunctus* and is an important site for lignocellulose digestion (Schwarz et al., 2023). Prior to dissection, we immobilized the specimens in a standard freezer at -20 °C for 60 minutes. In each case, the abdominal sclerites were removed to expose the intestine. The intestine was then sectioned into three parts (anterior, middle, and posterior) and placed on DNA/RNA (Zymo Research, Irvine, California, USA) and shipped to the Mikaelyan Lab at North Carolina State University (Raleigh, North Carolina, USA). The rest of the specimen was placed in 96% ethanol and kept as a voucher at the Colección Entomológica del Centro de Coleciones Biológicas (Universidad del Magdalena).

We combined the AHG of two specimens from the same colony to control for individual-level variation for DNA extraction (Supplementary Table S1). Microbial DNA was extracted using the Quick-DNATM Fecal/Soil Microbe Microprep Kit (Zymo Research) following the manufacturer’s protocol with slight modifications: (1) prior to the bead-beating step, we homogenized the guts with a sterile plastic mechanical pestles in DNA/RNA shield; (2) for the bead-beating step, the samples were further homogenized with the bead-beating solution using a Benchmark Beadblaster 24 homogenizer at 6.0 m/s in four cycles of 11s, with a 30s rest between cycles; and (3) we mixed 600 µL of the homogenized supernatant with the genomic lysis buffer instead of the 400 µL recommended by the manufacturer. We divided the microbial DNA extractions into three aliquots, one of which was kept as a voucher, and two were sent to Novogene (Beijing, China) for Bacterial and Archaeal metabarcoding. Bacterial ribosomal profiling was performed using the 16S V3-V4 region (341F-806r primer pair), and archaeal profiling was performed with the 16S V4-V5 region (Arch519F-Arch915R primer pair).

### Selection of Amplicon Sequence Variants and construction of phyloseq data sets

Demultiplexing and adapter trimming of raw sequencing reads was done by Novogene. We conducted subsequent analyses in R v 4.3.0 through RStudio v 2023.12.1.402 (Posit team, 2024) on servers hosted by the Minnesota Supercomputing Institute. Amplicon sequence variants (ASVs) were inferred using the DADA2 package (Callahan, McMurdie, et al., 2016), implemented within the Bioconductor Workflow for Microbiome Data Analyses (Callahan, Sankaran, et al., 2016), including quality filtering, dereplication, error-model learning, paired-end-merging, chimera removal, and taxonomic assignment.

First, individually for forward and reverse reads, we trimmed the reads by truncating at the first instance of a quality score less than 2; then we removed all reads containing Ns, those with an “expected error” higher than 2, and those with less than 200 nucleotides. We only kept the sequences for which both forward and reverse reads passed the trimming and filtering conditions. Subsequently, we deduplicated the forward and reverse reads. Afterwards, ASVs were inferred using the DADA2 method, which relies on a parameterized model of substitution error to distinguish sequencing errors from real biological variation (Callahan, Sankaran, et al., 2016). For this, we first estimated a separate error matrix for the deduplicated forward and reverse reads and subsequently inferred ASVs using the previously estimated error matrix. Finally, we merged the forward and reverse reads.

Next, we constructed a sequence table and inspected the distribution of lengths. After length inspection, we retained all merged sequences within a size range of 370-440 base pairs for the bacterial data set and 400-470 base pairs for the archaeal dataset. Later, we removed all chimeras using the removeBimeraDenovo function. Finally, we assigned taxonomy to the selected ASVs using the SILVA v138.2 (Quast et al., 2012) reference database. The sequence, taxonomy, and sample metadata tables were then consolidated into a phyloseq object.

We carried out the additional data handling and visualization using R package phyloseq (McMurdie & Holmes, 2013). To the data obtained from DADA2, we applied a taxonomic filter by removing eukaryotic and mitochondrial sequences. Furthermore, for the dedicated bacterial dataset, all archaeal ASVs were removed, and conversely, for the archaeal dataset, all bacterial ASVs were removed. Additionally, any ASVs that could not be classified at the phylum level were excluded. The resulting filtered ASV tables were utilized for alpha diversity analyses. An additional prevalence filter was applied to remove ASVs present in only a single sample. Subsequently, microbial phylogenetic trees were reconstructed using the ASVs retained after the prevalence filter following the phylogenetic tree reconstruction methods of the Bioconductor Workflow for Microbiome Data Analyses (Callaham et al. 2016) with one modification for the bacterial dataset: Due to the high number of bacterial ASVs, phylogenetic tree optimization was done using nearest neighbor interchange (NNI) rearrangement instead of the stochastic rearrangement setting. Finally, this prevalence-filtered ASV table was normalized by converting read counts to relative abundances. This normalized dataset was used for all subsequent analyses, including visualization of microbial composition, beta diversity, and phylosymbiosis.

### Measurements of Microbial Diversity

We estimated microbial alpha diversity on the taxonomically filtered data set using the Abundance-based Coverage Estimator (ACE) (Chao & Chiu, 2016) and Fisher’s alpha (FAD) (Fisher et al., 1943) diversity indices. Differences in alpha diversity across host genera and localities were tested using a one-way ANOVA test, and we measured the effect size of the overall difference using the R-squared (*r*²). Additionally, microbial Beta diversity was calculated from the normalized data set using Jaccard, Bray-Curtis, and weighted and unweighted UniFrac distances. Results were visualized using Principal Coordinate Analyses (PCoA).

### Detection of effects of host taxonomy and geographic distribution in microbial communities

To test the effects of phylogenetic relatedness and shared geographical distribution (i.e. locality) on microbiome composition and structure we performed permutational multivariate analysis of variance (PERMANOVA) using the adonis2 function of the R package Vegan (Oksanen et al., 2025) and Beta-dispersion analyses. Separate tests were run for each of the four distance matrices. To test the effect of locality we used a simple model (distmatrix ∼ locality). To test the effect of phylogenetic relatedness we used a sequential model (distmatrix ∼ tribe + genus + species). Sequential models estimate the significance of each term sequentially from first to last. Finally, to partition variance between phylogeny and locality, we used a marginal test model (distmatrix ∼ genus + locality). The marginal models estimate the unique variance explained by each factor after accounting for the other.

### Test of Phylosymbiosis

We tested the association between host phylogenetic relationships and host-associated bacterial and archaeal community structure following the practical framework of Lim & Bordenstein (2020). A host phylogenetic tree was reconstructed under the Maximum Likelihood criterion using IQ-TREE2 (Minh et al., 2020) based on a concatenated alignment of one nuclear (WGS) and two mitochondrial (12S and COXI) genes. For each of the four beta-diversity distance matrices we constructed a microbiome dendrogram (phylobiome) using hierarchical clustering with the UPGMA method using the hclust function in R.

To assess whether a phylobiome recapitulated the host phylogeny better than expected by chance, we performed a topology-based randomization test. First, we calculated the topological distance between each phylobiome and the true host tree using the Robinson-Foulds distance and the Matching Cluster distance using TreeCmp (Goluch et al., 2020). We then generated a null distribution by calculating the same distance metrics between the phylobiome and 1,000 randomly generated host phylogenetic trees. Subsequently a one-tailed *p*-value was derived as the proportion of random trees that had a distance metric value lower or equal to the observed value between the phylobiome and the host tree. A *p*-value < 0.05 was considered evidence of significant phylosymbiosis for that phylobiome.

## Results

### Microbiome Composition and Structure

For the bacterial data set, we successfully obtained microbial profiles for 31 samples, with an average input read count of 204,820 (+-10,436). After processing with the DADA2 pipeline, the samples had an average read count of 141,610 (+-17,852) (Supplementary Table S2). For Archaea, we obtained microbial profiles for 28 samples, with an average input read count of 190,424 (+-31,620), and an average read count of 97,171 (+-23,560) following read processing (Supplementary Table S3).

Following the application of the taxonomic filter, we obtained 31,226 and 2,102 ASVs for Bacteria and Archaea, respectively. The ANOVA test showed a significant effect of host genera on ASV counts for both Bacteria and Archaea (*p* < 0.05) in both alpha diversity indices (Fig. 2 A, C). However, the overall effect size for Bacteria (*r*^2^ = 0.593 for ACE and *r*^2^ = 0.706 for FAD) was much higher than that for Archaea (*r*^2^ = 0.366 for ACE and *r*^2^ = 0.277 for FAD). This indicates that host genera identity accounts for a larger proportion of the variance in bacterial ASV richness compared to archaeal ASV richness. In contrast, no significant differences were found between the different localities in both microbial profiles (Fig 2 B, D).

**Figure 2.**
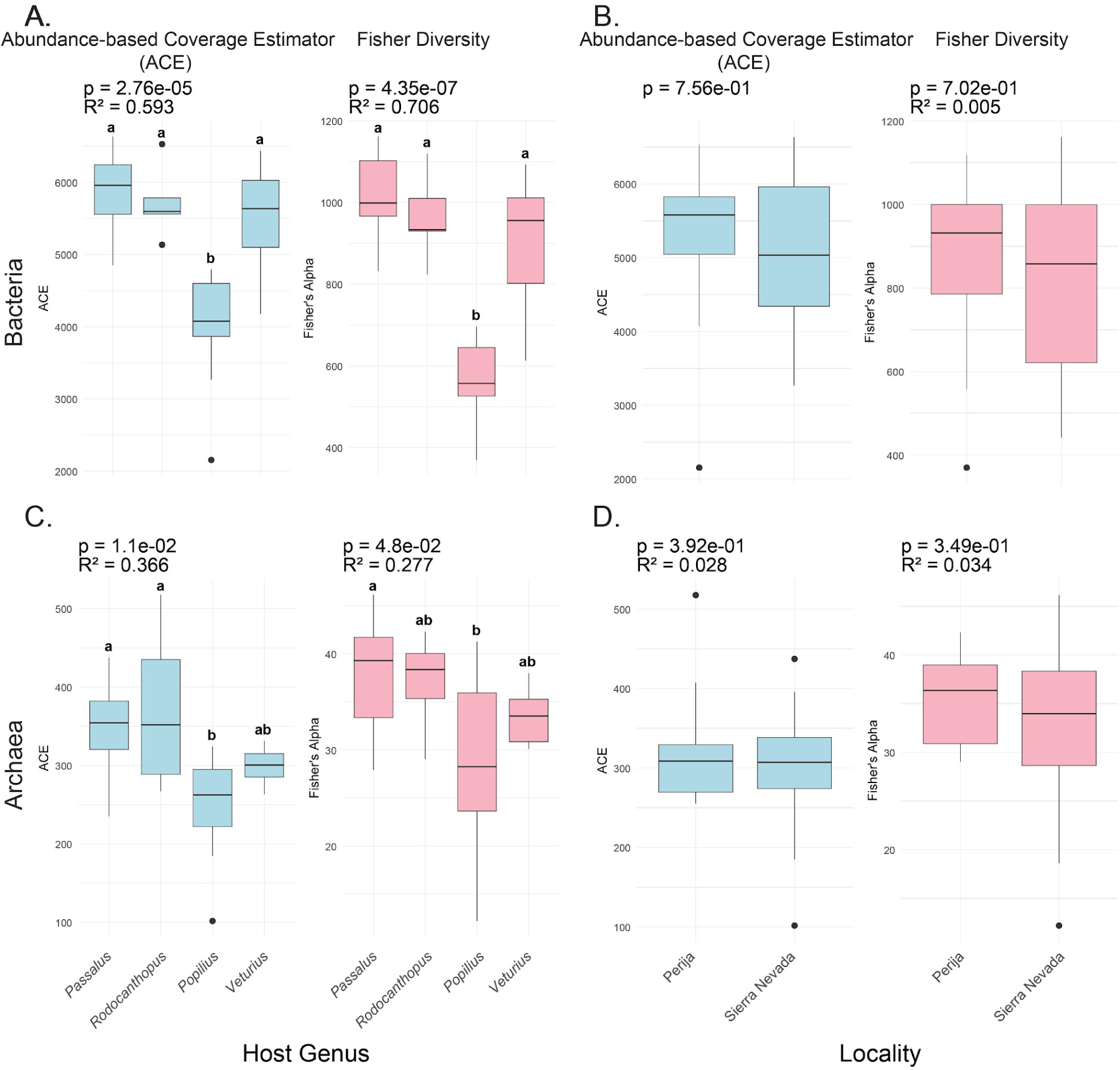
Alpha diversity of bacterial and archaeal gut communities across Passalidae species. ACE and Fisher indices calculated from taxonomically filtered ASVs. Boxes represent genus-level (A, C) or locality (B, D) distributions. Significant differences among genera were detected from one-way Anova test, (R²) effect size of the overall difference. A) Host genera, Bacteria; B) Locality, Bacteria; C) Host genera, Archaea; D) Locality, Archaea.

Our microbial profiling results show a difference in the taxonomic knowledge gaps of bacteria and archaea inhabiting the gut of Passalidae. For Bacteria, we were able to classify 191 family level groups and 291 generic level groups corresponding to 89.6 and 49.5% of reads respectively. For Archaea, we were able to classify 7 family level groups and 7 generic level groups corresponding to 99.7 and 97.29% of archaeal reads respectively. For Bacteria, 53% of the recovered ASVs, representing 50.4% of total reads, could not be classified to genus level. In contrast, only 19% of Archaea ASVs, representing only 2.71% of the total reads, were not classified at the genus level. This pattern is kept across all taxonomic rankings (Supplementary Table S4).

We identified the top 20 classified families and genera for Bacteria (Fig. 3); collectively they accounted for 79.86 ± 3.12% and 39.10 ±3.25% of the bacterial community, respectively (Fig. 3A, B). For all samples Christensenellaceae was the most abundant family; while Christensenellaceae R-7 group was the most abundant genus level group. For Archaea we were able to name 6 family and genus level groups (Fig. 4); together they represented 99.96 ± 0.03% and 96.95 ± 2.24% of the archaeal community, respectively (Fig. 4A, B). Methanosarcinaceae was the most abundant family and Methanimicrococcus the most abundant genus in all samples.

**Figure 3.**
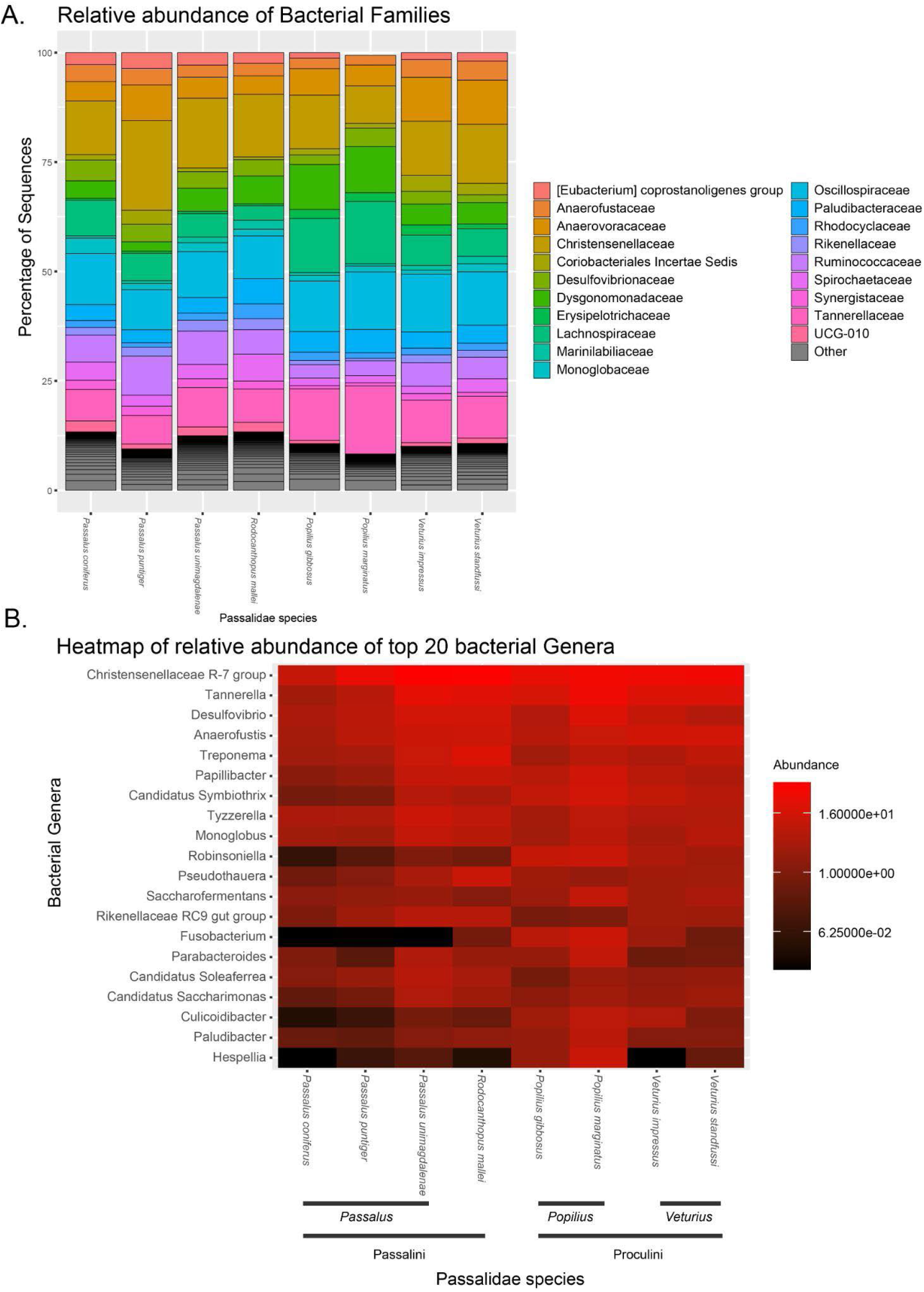
Bacterial gut microbiome composition across Passalidae species. A) Relative abundance of the top 20 bacterial families detected in the anterior gut. Bars represent species-level means; colors indicate families. B) Heatmap of the top 20 genera across species, showing compositional differences among Passalidae genera and highlighting dominant groups such as Treponema and Christensenellaceae R-7.

**Figure 4.**
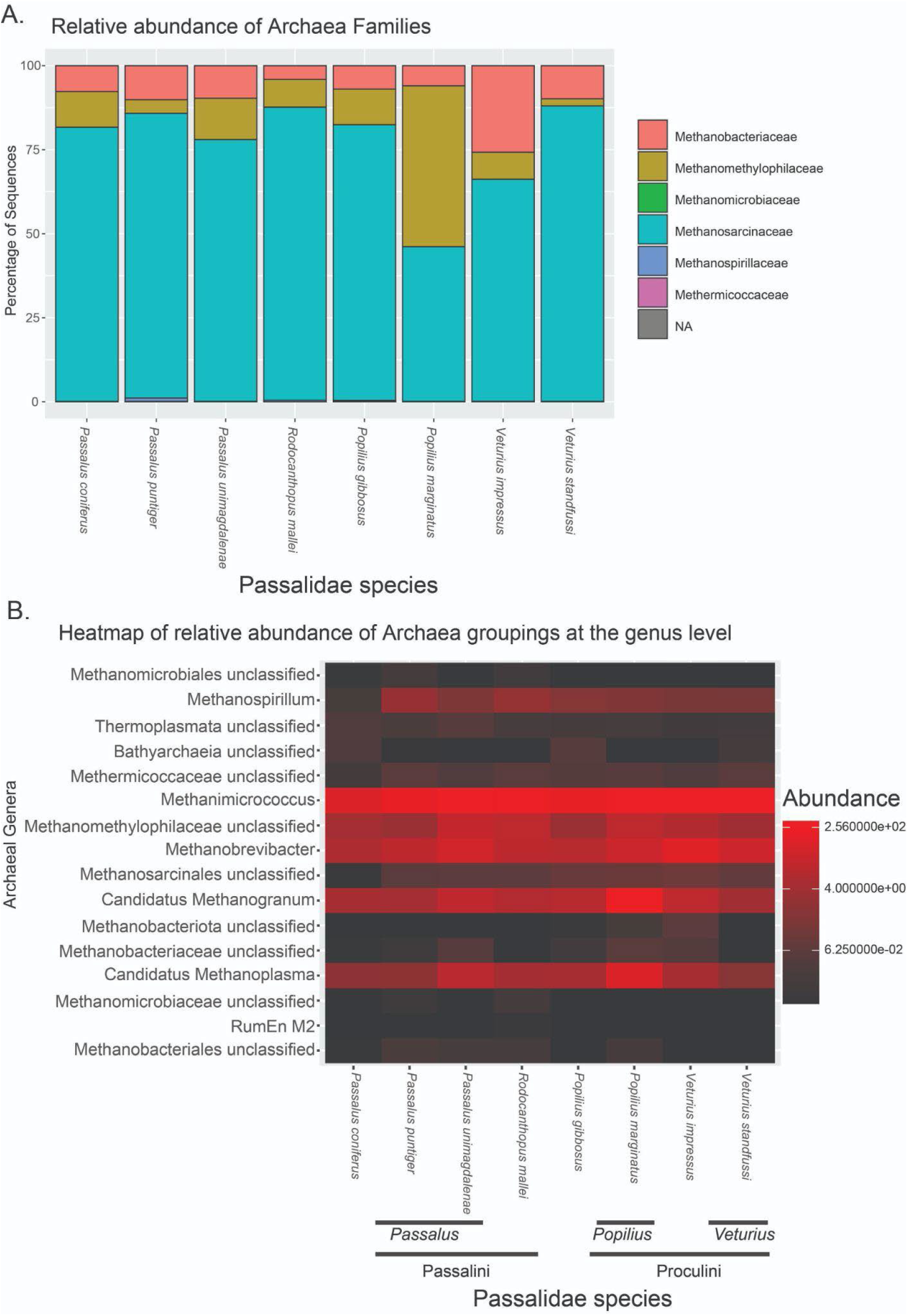
Archaeal gut microbiome composition across Passalidae species. A) Relative abundance of archaeal families detected in the anterior gut. Bars represent species-level means; colors indicate families. B) Heatmap of the genera across species.

Additionally for Bacteria, in the examination of the top 20 most abundant ASVs we can identify three distinct dominance patterns (Figure 5). Species in the tribe Passalini show high abundance of Anaerofustis, and a set of five ASVs classified to the Christensenellaceae R-7 genus level group. *Popilius* have a high abundance of Lachnospiraceae CHKCI001, Papilibacter, Fusobacterium, and one ASV classified to Christensenellaceae R-7 group, different from those that are abundant in Passalini. Finally, *Veturius* shows a differential high abundance of an ASV classified as an unidentified Tannerellaceae, and another unclassified Anaerovoraceaea. This compositional dissimilarity can be detected in the beta diversity PCoA plots under all indices. For Archaea a similar but attenuated pattern is shown (Supplementary Fig. S1).

**Figure 5.**
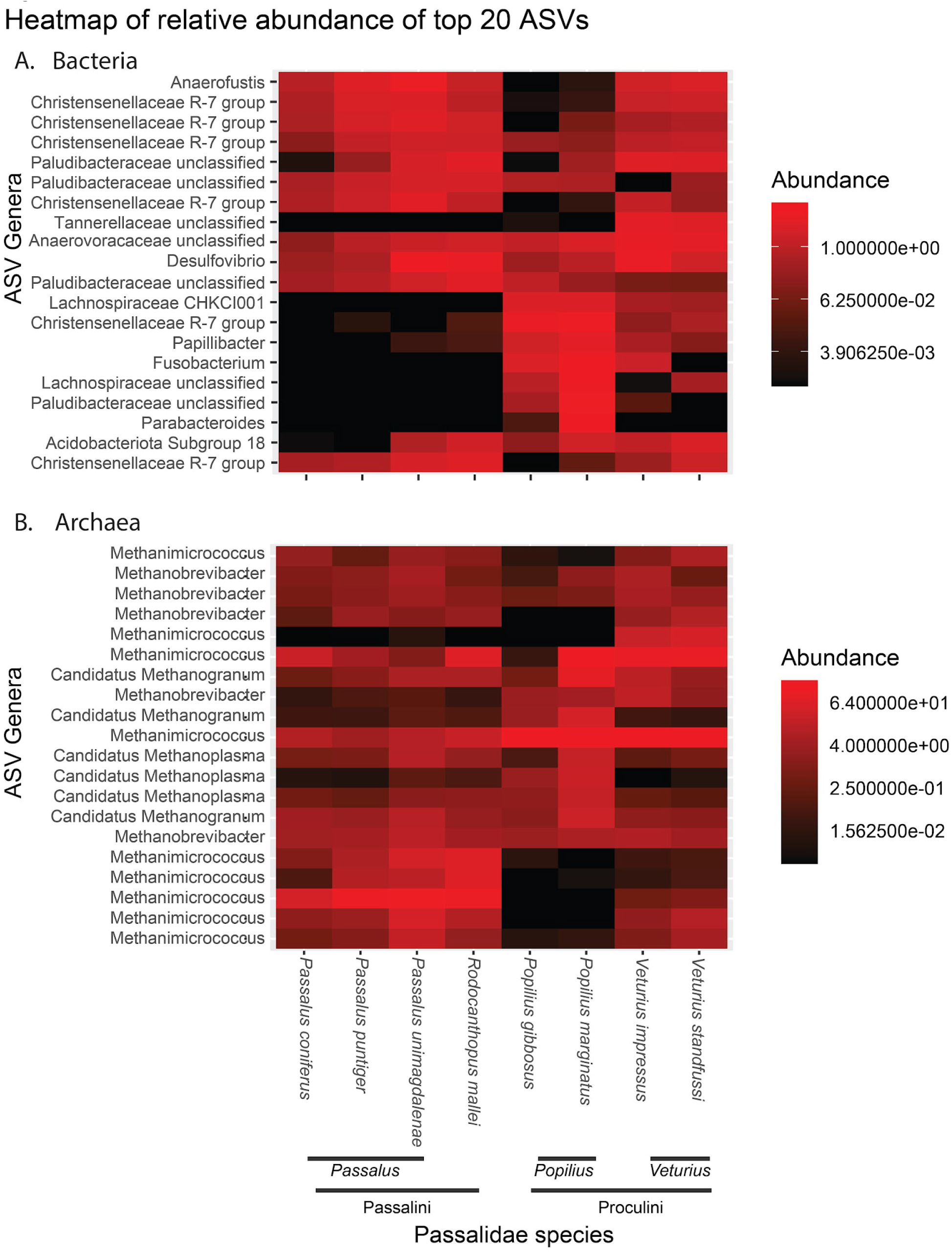
Heatmap of dominant bacterial and archaeal taxa in the anterior hindgut of Passalidae. Heatmap showing relative abundances of the top 20 most abundant ASVs across all samples. Columns represent species means. Rows represent taxa scaled by row-wise abundance (Z-scores). A) Bacteria; B) Archaea.

### Effects of host taxonomy and geographic distribution in microbial communities

In general, our analyses show a larger effect of host taxonomy vs geographical distribution in the microbial communities of bess beetle anterior guts. While we detect a structure of microbial diversity dependent on host taxonomy, the PCoA analyses show no pattern regarding sample localities in both Bacteria (Fig. 6) and Archaea (Fig. 7). For Bacteria, our sequential model to test for the effects of host taxonomy was statistically significant (*p* < 0.05) in almost all taxonomic levels and beta diversity indices (Table 2); the only exception being at the host species level under the weighted UniFrac index. However, Dispersion differed significantly among groups at the tribe and species levels for Jaccard, Bray-Curtis, and unweighted UniFrac (Supplementary Table S5). Meanwhile, the locality only model was statistically significant for all indices except both UniFrac metrics; nonetheless, *r*^2^ values were low in all cases (5-8 %). A similar pattern was observed in our marginal model that included host taxonomy (genus level) and locality, where genus was statistically significant for all metrics and locality was statistically significant for three of the four with weighted UniFrac being the exception.

**Figure 6.**
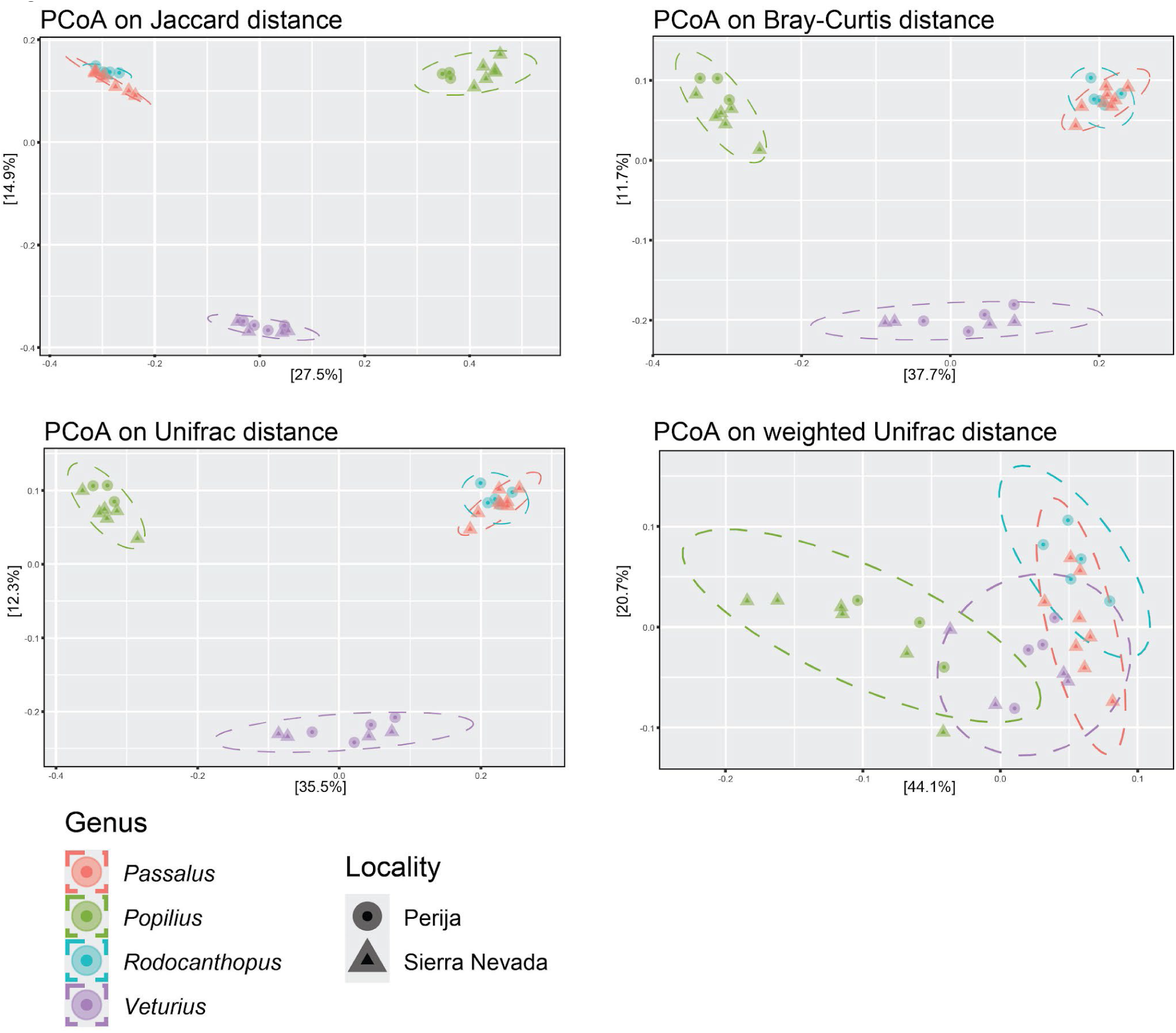
Beta diversity of passalid gut bacterial microbiomes shows clustering by host genus. Principal Coordinates Analysis (PCoA) based on Jaccard, Bray-Curtis, unweighted UniFrac, and weighted UniFrac distance matrices. Points represent extraction samples; colors indicate host genus and shapes indicate locality. Across all metrics, samples cluster primarily by genus rather than locality, suggesting host phylogeny plays a stronger role than geography in structuring gut bacterial communities.

**Figure 7.**
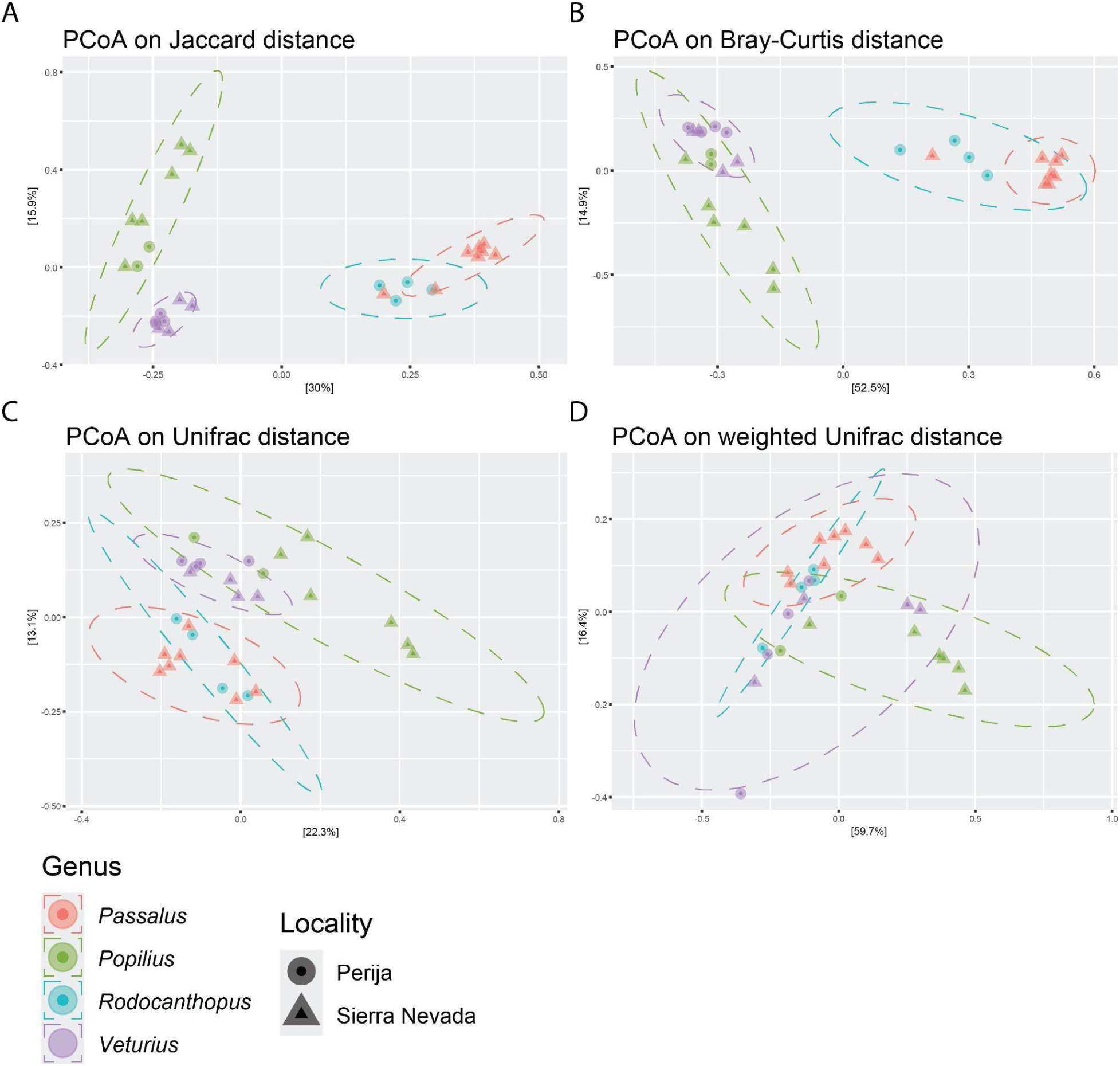
Beta diversity of passalid gut archaeal microbiomes shows clustering by host genus. Principal Coordinates Analysis (PCoA) based on Jaccard, Bray-Curtis, unweighted UniFrac, and weighted UniFrac distance matrices. Points represent extraction samples; colors indicate host genus and shapes indicate locality. Across most metrics, samples cluster primarily by genus rather than locality, suggesting host phylogeny plays a stronger role than geography in structuring gut archaeal communities. This pattern was not recovered for the weighted UniFrac distance matrix.

**Table 2.** Results of permutational multivariate analysis of variance (PERMANOVA) using the adonis2 of beta diversity matrices of bacterial microbiome using 3 models: sequential (dist ∼ Tribe + Genus + Species), single (dist ∼ locality), and marginal (dist ∼ locality + genus) tests.

|  |  | Jaccard |  |  | Bray-Curtis |  |  | Unifrac |  |  | Weighted Unifrac |  |  |
|---|---|---|---|---|---|---|---|---|---|---|---|---|---|
| Model |  | R2 | F | Pr | R2 | F | Pr | R2 | F | Pr | R2 | F | Pr |
| dist ~ Tribe + Genus + Species, sequential test | Tribe | 0.24 | 19.51 | **0.0010** | 0.24 | 19.247 | **0.0010** | 0.29 | 20.06 | **0.0010** | 0.26 | 14.82 | **0.0010** |
|  | Genus | 0.25 | 10.24 | **0.0010** | 0.26 | 10.608 | **0.0010** | 0.24 | 8.19 | **0.0010** | 0.25 | 7.23 | **0.0010** |
|  | Species | 0.24 | 4.89 | **0.0010** | 0.22 | 4.444 | **0.0010** | 0.16 | 2.74 | **0.0010** | 0.10 | 1.49 | 0.1039 |
| dist ~ locality | Locality | 0.08 | 2.49 | **0.0180** | 0.08 | 2.3522 | **0.0220** | 0.05 | 1.51 | 0.1409 | 0.05 | 1.54 | 0.1449 |
| dist ~ locality + genus, marginal test | Genus | 0.49 | 9.62 | **0.0010** | 0.50 | 9.93 | **0.0010** | 0.52 | 10.3 | **0.0010** | 0.49 | 8.89 | **0.0010** |
|  | Locality | 0.08 | 4.89 | **0.0010** | 0.07 | 4.44 | **0.0010** | 0.05 | 2.99 | **0.0130** | 0.03 | 1.62 | 0.1299 |

**Table 3.**
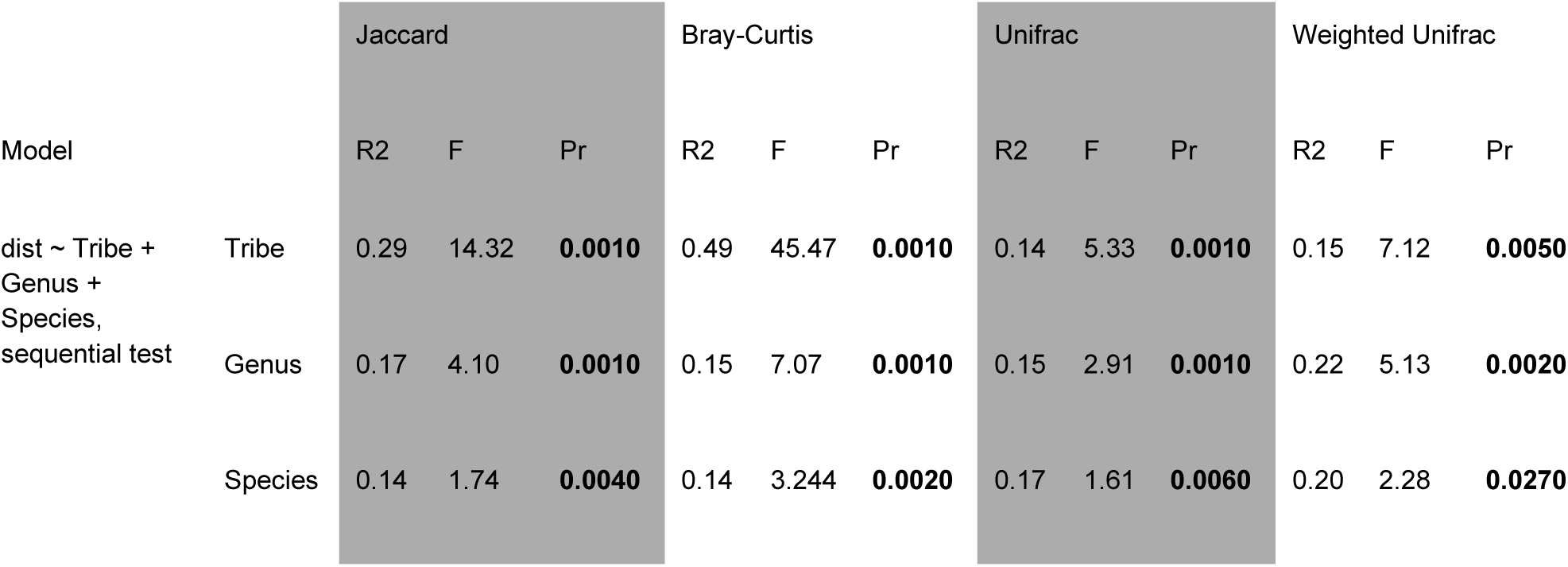
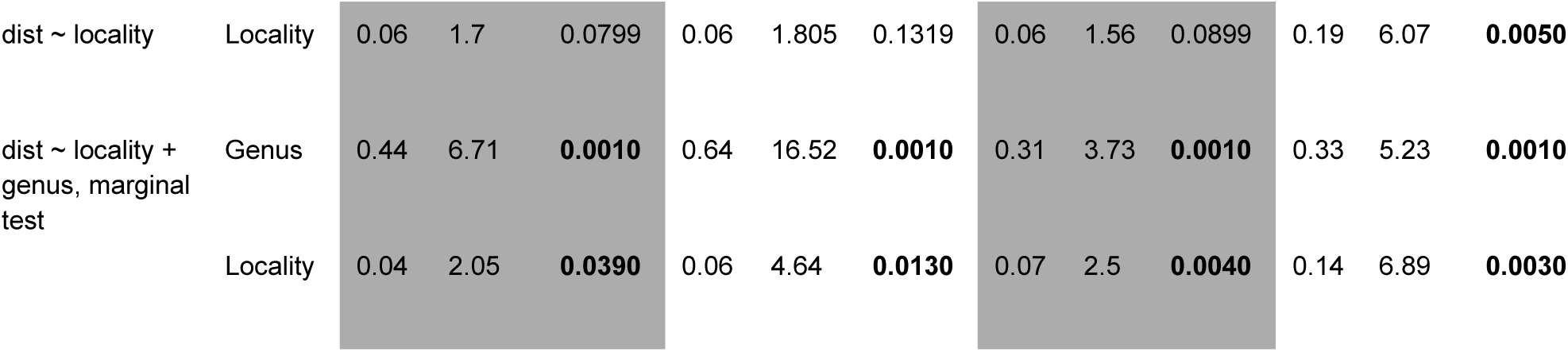
Results of permutational multivariate analysis of variance (PERMANOVA) using the adonis2 of beta diversity matrices of archaeal microbiome using 3 models: sequential (dist ∼ Tribe + Genus + Species), single (dist ∼ locality), and marginal (dist ∼ locality + genus) tests.

|  |  | Jaccard |  |  | Bray-Curtis |  |  | Unifrac |  |  | Weighted Unifrac |  |  |
|---|---|---|---|---|---|---|---|---|---|---|---|---|---|
| Model |  | R2 | F | Pr | R2 | F | Pr | R2 | F | Pr | R2 | F | Pr |
| dist ~ Tribe + Genus + Species, sequential test | Tribe | 0.29 | 14.32 | **0.0010** | 0.49 | 45.47 | **0.0010** | 0.14 | 5.33 | **0.0010** | 0.15 | 7.12 | **0.0050** |
|  | Genus | 0.17 | 4.10 | **0.0010** | 0.15 | 7.07 | **0.0010** | 0.15 | 2.91 | **0.0010** | 0.22 | 5.13 | **0.0020** |
|  | Species | 0.14 | 1.74 | **0.0040** | 0.14 | 3.244 | **0.0020** | 0.17 | 1.61 | **0.0060** | 0.20 | 2.28 | **0.0270** |
| dist ~ locality | Locality | 0.06 | 1.7 | 0.0799 | 0.06 | 1.805 | 0.1319 | 0.06 | 1.56 | 0.0899 | 0.19 | 6.07 | **0.0050** |
| dist ~ locality + genus, marginal test | Genus | 0.44 | 6.71 | **0.0010** | 0.64 | 16.52 | **0.0010** | 0.31 | 3.73 | **0.0010** | 0.33 | 5.23 | **0.0010** |
|  | Locality | 0.04 | 2.05 | **0.0390** | 0.06 | 4.64 | **0.0130** | 0.07 | 2.5 | **0.0040** | 0.14 | 6.89 | **0.0030** |

For Archaea, similar results to the bacterial communities were observed when testing effects of host taxonomy, with the difference that host species had a statistically significant effect under the weighted UniFrac index. In the case of the locality only model, specimen geographic distribution had no effect in the archaeal communities except for the weighted UniFrac index. Likewise, to the bacterial results, the marginal model which combined host taxonomy and locality, showed a significant effect of both predictors but with a greater effect of host taxonomy. Different from Bacteria, the beta dispersion analyses in Archaea were significant in all variables for the Jaccard and unweighted UniFrac indices, and not significant for Bray-Curtis and Weighted UniFrac, suggesting the effects shown in our PERMANOVA are more robust for archaeal community structure than community composition.

### Phylosymbioses

For Bacteria, the Jaccard, Bray-Curtis and unweighted UniFrac distance matrices produced the same phylobiome (Fig 8B), while weighted UniFrac resulted in a different phylobiome where the position of *Passalus punctiger* was different (Fig 8C). In contrast, for Archaea, each distance matrix resulted in a different phylobiome (Fig. 9B-E).

**Figure 8.**
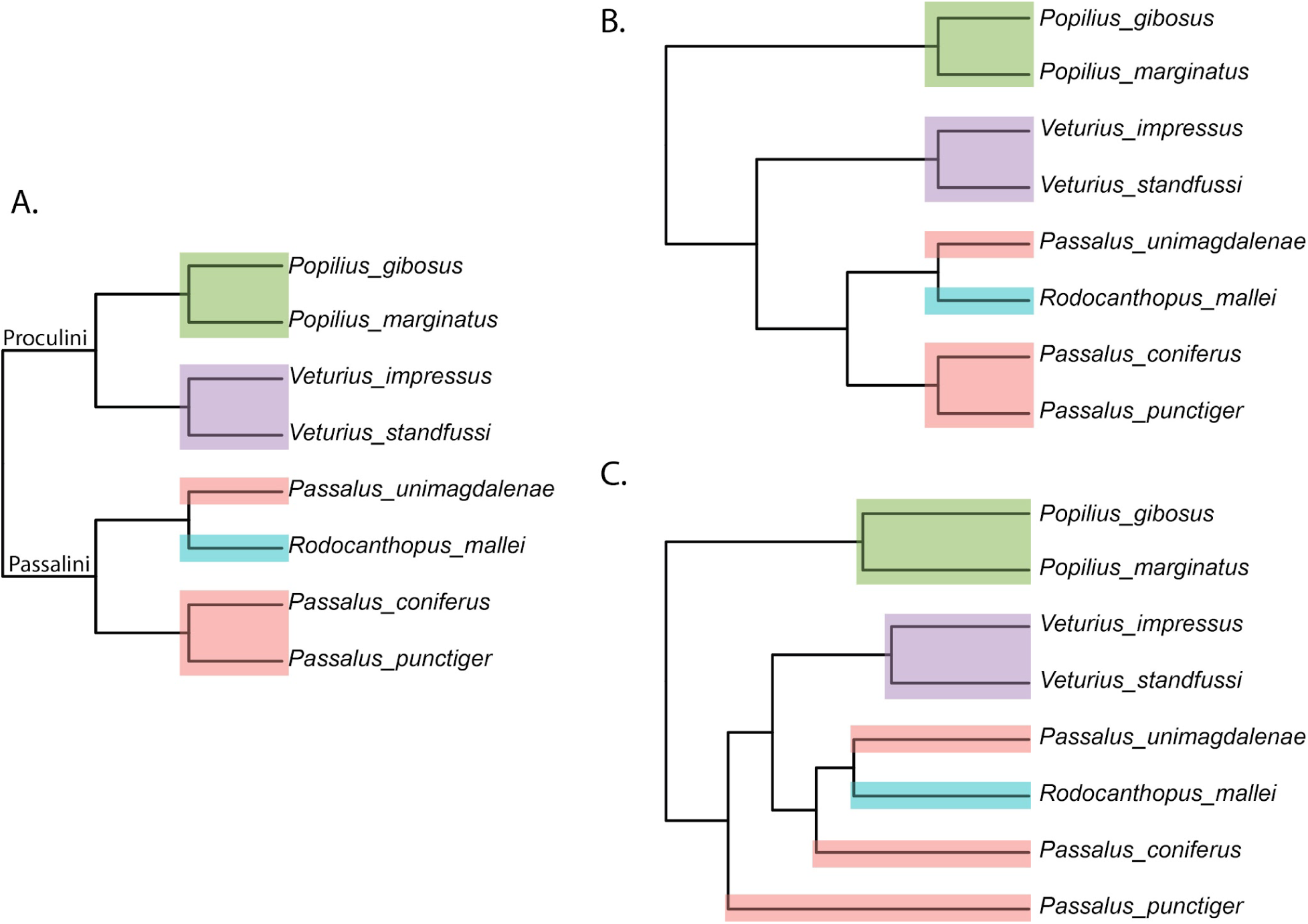
Host phylogeny and bacterial microbiome dendrograms reveal phylosymbiosis in Passalidae. (A) Maximum-likelihood phylogeny of Passalidae species included in this study, shown here for comparison with microbiome clustering. (B, C) UPGMA dendrograms of gut bacterial communities based on B) Jaccard, Bray-Curtis, unweighted UniFrac, and C) weighted UniFrac distance matrices. Tip colors correspond to the host genera.

**Figure 9.**
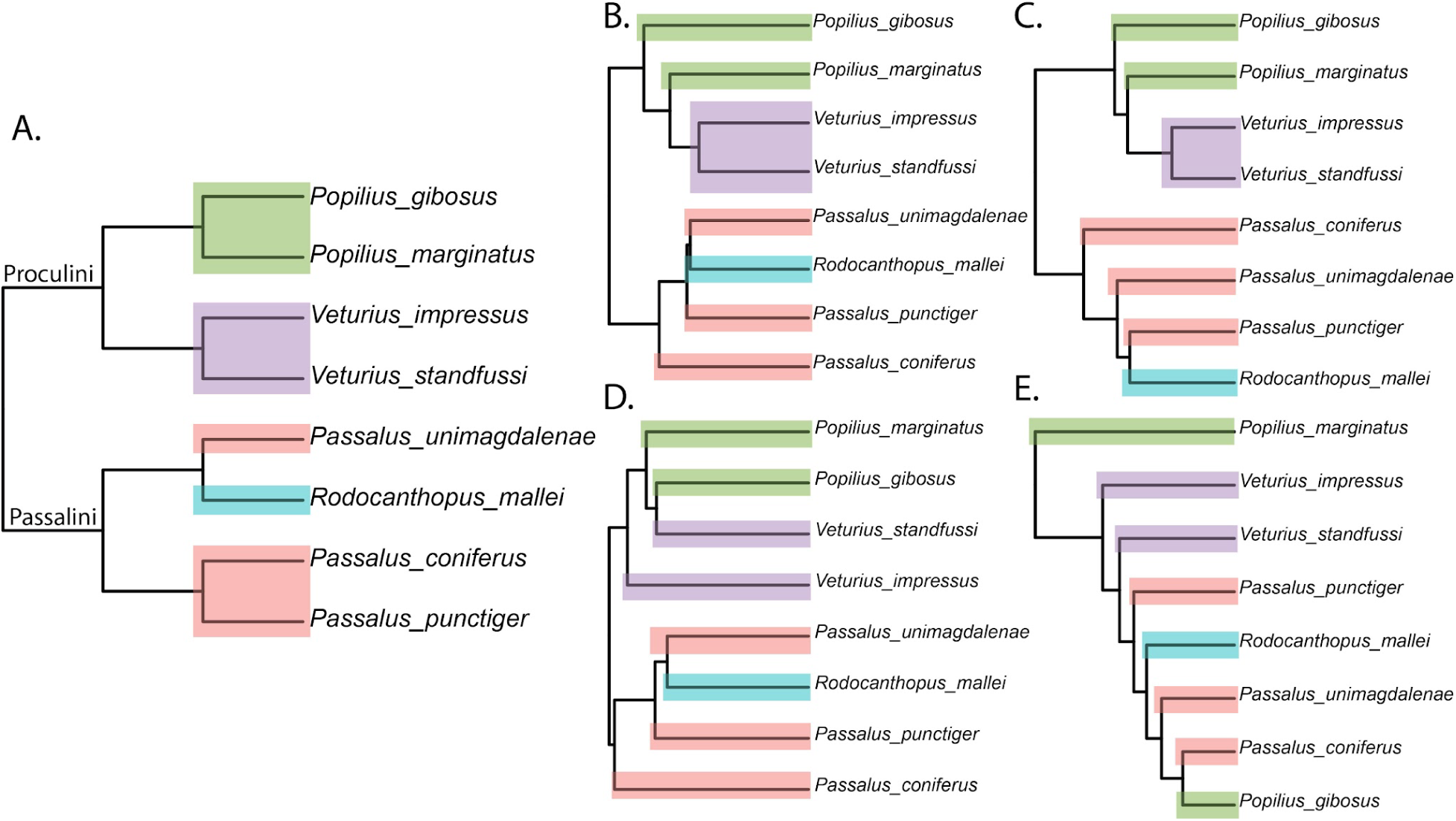
Figure 8. Host phylogeny and archaeal microbiome dendrograms reveal phylosymbiosis in Passalidae. (A) Maximum-likelihood phylogeny of Passalidae species included in this study, shown here for comparison with microbiome clustering. (B-E) UPGMA dendrograms of gut bacterial communities based on B) Jaccard; C. Bray-Curtis; D) unweighted UniFrac; and E) weighted UniFrac distance matrices. Tip colors correspond to the host genera.

The results of the topology-based randomization tests indicate that most of the resulting phylobiomes resemble host phylogeny more than expected by chance (Table 4). In Bacteria, the only exception was the weighted UniFrac distance phylobiome under the matching cluster criteria, where 6% of random trees had a distance metric value lower or equal to the observed value between the phylobiome and the host tree. Meanwhile in Archaea, the phylobiome produced by weighted UniFrac had 100% random trees with a distance metric value lower or equal to the observed value between the phylobiome and the host tree using both tree matching criteria.

**Table 4.** Summary of one tailed randomization test, values in columns represent the proportion of 1000 randomly generated phylogenetic trees with a smaller phylogenetic distance to the phylobiome than that between the phylobiome and the host tree.

|  | Bacteria |  | Archaea |  |
|---|---|---|---|---|
| Matrix | RF | MC | RF | MC |
| jc | **0** | **0.003** | **0.003** | **0.005** |
| bc | **0** | **0.003** | **0** | **0.005** |
| Uni | **0** | **0.003** | **0.003** | **0.005** |
| wUni | **0.002** | 0.067 | 1 | 1 |

On one hand, the bacterial phylobiomes maintain the relationships between closely related species. However, they fail to recover the phylogenetic structure at the host tribe level. On the other hand, in Archaea all phylobiomes recover the host tribe level phylogenetic relationships but fail to recover similarity patterns that resemble the phylogenetic relationships between closely related host species.

## Discussion

By sampling multiple passalid species across two geographically close but biogeographically distinct Colombian mountain systems, pairing host phylogenies with bacterial and archaeal ribosomal profiles, and testing for tree congruence, we provide the first comparative and phylogenetically informed assessment of passalid gut microbiota. Our results reveal three main patterns: (i) substantial bacterial taxonomic novelty and deep under-classification in the anterior hindgut microbiota of Passalidae contrasted with comparatively low-diversity and taxonomically well-resolved archaeal assemblages, (ii) strong and statistically significant host taxonomy effects on alpha and beta microbial diversity; compared to modest locality effects, limited to some beta diversity metrics, (iii) clear evidence of bacterial and archaeal phylosymbiosis at different host phylogenetic levels. These results suggest that the microbial communities in the anterior hindguts of Passalidae are correlated more with the evolutionary history of the group than the occupied geographic area. Together, these patterns position Passalidae as a major system for dissecting how gut compartmentalization, social transmission, and ecological filtering interact to shape lignocellulose-feeding symbioses.

### Microbiota composition and novel microbial communities

A defining feature of the passalid gut microbiota is the extent of bacterial under-classification. Similar results of taxonomic underclassification of bacteria found in this study have been reported for a large number of wood feeding insects (Beza-Beza et al., 2024). This highlights the need for further work both in increasing the number of sequences of insect associated microbes in databases and classifying more insect-associated bacteria. Wood-feeding insects repeatedly evolve gut symbioses that are taxonomically novel, highly structured, and tightly coupled to host physiology and evolutionary history (Beza-Beza et al., 2024; Mikaelyan, Dietrich, et al., 2015). Termites provide the best-known example, but passalid beetles represent an independent radiation of subsocial saproxylophagous insects whose digestive biology and social ecology make a strong a priori case for specialized gut consortia.

Deep microbial under-classification in Passalidae parallels the taxonomic knowledge gaps first described in termites (Supplementary Table S4). In multiple microbial surveys of termites, a large fraction of 16S reads could not be assigned below very high taxonomic ranks, prompting the creation of termite-specific databases (e.g., Mikaelyan, Köhler, et al., 2015) and eventually reference-independent genome reconstruction (Hervé et al., 2020; Manjula et al., 2015). From 30 higher-termite gut metagenomes, Hervé et al. (2020) recovered 589 MAGs, anchoring 16S-only “termite-group” clades such as TG3 within Fibrobacteres, expanding *Treponema* I diversity, and revealing termite-specific Bathyarchaeota lineages. These MAGs fundamentally reshaped interpretations of termite gut structure and function and exposed the inadequacy of general 16S databases for insect-microbe systems dominated by deeply host-restricted lineages.

Our passalid dataset shows a similar pattern: nearly half of bacterial reads could not be assigned to genus. Given that termite-specific clades remained “unclassified” until genome-resolved analyses were available, a substantial portion of this unclassified bacterial fraction in the passalid microbiota may represent host-associated lineages that are poorly represented in current databases. However, unlike termites, passalids have not yet been surveyed with MAG-rich phylogenomics, so we cannot yet say whether they harbor beetle-specific phylum- or class-level clades analogous to termite “TG” lineages (Brune & Dietrich, 2015) or instead represent somewhat shallower radiations within known phyla. Alternatively, MAG-rich phylogenomic sampling could increase our classification capabilities resulting in a group of already described but poorly represented lineages in 16S bacterial data bases.

Apart from *Odontotaenius disjunctus*, passalid microbiota evolution has largely remained a major blind spot in the study of wood-feeding insects. Thus, our results also need to be interpreted in light of what is already known about microhabitat structure and fibrolytic guilds in this species. (Schwarz et al., 2023) showed that the anterior hindgut (AHG) is a key site of lignocellulose digestion in *O. disjunctus*, harboring the bulk of gut bacterial biomass and exhibiting high cellulase and xylanase activities. Using a Percoll-based protocol, they enriched ingested wood fibers from AHG lumen and demonstrated that these particles host a distinct fiber-associated microbiota, compositionally separated from fiber-free lumen communities and strongly enriched in insect-restricted lineages of *Lactococcus*, *Turicibacter*, *Terponema*, and the Christensenellaceae R7 group. Wood particles contributed a large share of total hydrolytic activity, and the authors argued that wood fibers constitute a crucial microhabitat for cellulose- and xylan-degrading bacteria in the passalid hindgut.

Several of the genera that define this fiber-associated guild occur across the Colombian passalid species we sampled (Fig 3). For example, we recovered a high abundance of the Christensenellaceae R7 group, with 6 of our top 20 most abundant ASVs belonging to this group (Fig. 5). Additionally, we also identified *Treponema* as one of the top 20 genera within our study (Fig. 3B). Even though our design did not fractionate fiber-attached versus planktonic communities, the anterior hindgut samples inevitably integrate both microhabitats. The recurrence of these taxa across species suggests that the fibrolytic strategy resolved at high spatial resolution in *O. disjunctus* may not be unique to that species, but may instead represent a broader feature of the passalid anterior hindgut. This interpretation is consistent with termite work showing that fiber-attached consortia and lumen communities are compositionally distinct but functionally overlapping, with both fractions contributing to lignocellulose depolymerization (Mikaelyan et al., 2014; Tokuda et al., 2018).

A key unresolved question is whether the fiber-associated guilds described in *O. disjunctus* represent a passalid-specific innovation or specialized form a broader scarabaeoid trait. No study has yet compared passalid gut communities with those of their closest relatives (e.g., geotrupids, trogids, lucanids, scarab beetles; Beza-Beza et al., 2024), some of which are considered saproxylophagous, using equivalent sampling of hindgut compartments or particle-attached fractions. Existing microbiota surveys of these sister groups – typically based on whole-gut homogenates – report communities dominated by generalist fermenters (e.g., Enterobacteriaceae, Lactobacillales, Clostridia), but they do not resolve whether analogous fiber-associated guilds exist, nor whether these lineages possess microhabitats comparable to the hypertrophied, microanoxic anterior hindgut of passalids. A phylogenetically controlled comparison therefore remains an important next step. Testing whether the fiber-attached consortia (comprised of *Lactococcus*, *Turicibacter*, *Treponema*, and Christensenellaceae R7) are unique to Passalidae, or instead represent a deeper scarabaeoid condition, would clarify whether the anterior-hindgut fibrolytic complex evolved within Passalidae or was co-opted and elaborated from ancestral scarab beetle states. Our results highlight this comparative gap and provide a framework for future work targeting sister-lineage contrasts.

### Effects of host taxonomy and geographical distribution on alpha and beta diversity of microbial communities

Alpha diversity analyses of this study found a strong effect of host taxonomy and no effect of locality on microbial richness. Across other insects with wood-feeding species, bacterial alpha diversity is better predicted by microhabitat structure than by diet stage or social behavior per spare. On one hand, species of the family Termitidae feeding on soil, humus, or grass often harbor more diverse bacterial communities than strict wood feeders, but diet category alone is unlikely to explain the patterns observed here because the passalid species included in this study share broadly similar wood-feeding and wood-dwelling habits. Instead, unmeasured host-associated factors like gut pH, oxygen penetration, and H₂ gradients across compartments could explain a substantial portion of microbial alpha diversity patterns (Köhler et al., 2012; Mikaelyan et al., 2017). Principles similar to these hold in wood-feeding cockroaches and passalids: Panesthiine cockroaches exhibit anoxic centers and steep redox gradients across compartments, with the hindgut hosting the most diverse, dense communities (Bauer et al., 2015), while in *O. disjunctus* all compartments possess anaerobic cores but the AHG concentrates bacterial load and fermentation (Ceja-Navarro et al., 2019). In our multi-species dataset, bacterial diversity differed among host genera but showed low within-lineage dispersion, consistent with strong anatomical and physicochemical constraints within clades and relatively homogenized communities within species. Social transmission in subsocial insects typically narrows rather than inflates interindividual variability – burying beetle larvae that receive parental care, for example, acquire a gut community dominated by maternal taxa, whereas no-care larvae are colonized by diverse carcass-derived bacteria (Wang & Rozen, 2017). The low within-species β-dispersion we observe in passalids is consistent with a similar reduction in stochastic colonization via frass provisioning and gallery reuse.

At the community level, β-diversity in our passalid dataset is structured more strongly by host lineage than by geographic distribution. However, whether this pattern reflects phylogenetic niche conservatism in the microbiota remains to be determined. For example, in Bacteria, the sequential model designed to test phylogenetic effect was significant for almost all taxonomic levels, with the exception of species for the weighted UniFrac distances (Table 2). Nonetheless, tests of overdispersion indicated that the values associated with different tribes and species were significantly different. Locality also had significant effects for the Jaccard and Bray-Curtis distance matrices but only explained a small proportion of the variance. Thus, bacterial community structure appears to carry a strong host-taxonomic signal, especially at the genus level, while geography contributes a secondary and comparatively weak component.

In Archaea, composition (Fig. 9 A, C) and structure (Fig. 9 B, D) are significantly influenced by host taxonomy at all levels. However, measures of composition alone (Jaccard, and unweighted UniFrac) show overdispersion for all factors tested, violating the assumptions of our models. In contrast, measures that included structure (Bray-Curtis, and weighted UniFrac) did not show significant differences of dispersion among the factors, indicating that when structure is taken into account we observe a strong effect of host lineage, accounting for approximately 78% of the differences observed by Bray-Curtis, and 57% of the differences observed in the weighted UniFrac distance matrix. Locality shows no significant effect on the Bray-Curtis distance matrix, and significant effect for weighted UniFrac explaining 19% of the differences when tested by itself and 14% when tested alongside the genus. This suggests that archaeal communities may be more broadly shared across passalid genera than bacterial communities, but that their dominant lineages vary in abundance in ways that retain host-associated structure.

The patterns observed in our study parallel reports for other decomposer insects. Neotropical dung beetles, host tribe explains more variation in gut communities than habitat or food preference (Jácome-Hernández et al., 2023), and in bark beetles, core bacteriomes persist across multiple populations with little signal of geographic distance (Hernández-García et al., 2018). Termite surveys across continents likewise find that genus-level clustering is robust and monophyletic for bacteria under unweighted UniFrac, with diet mostly shifting relative abundances rather than core membership (Abdul Rahman et al., 2015; Mikaelyan, Dietrich, et al., 2015). Our results are congruent with this macroecological structure: host taxonomy explains more variation than locality, while geographic distribution adds a smaller but detectable component. Such patterns are consistent with the passalids possessing shared adaptations in gut architecture, intestinal physicochemistry, and social transmission. As has been suggested for termites (Brune & Dietrich, 2015; Chakraborty et al., 2023; Huang et al., 2016; Su et al., 2016), such adaptations likely generate lineage-specific assembly rules in passalids, while local environmental factors (wood chemistry, temperature, microbial pool) modulate community composition without erasing host signals.

In contrast to the bacteria, the archaeal component of passalid gut microbiotas in our study was low in diversity and taxonomic breadth and showed significant structuring at different host phylogenetic levels (e.g., tribal for Archaea, and genus level groups for Bacteria). This domain-level asymmetry is expected because methanogens form narrow functional guilds constrained by H₂ availability, anoxia, and extreme compartmental pH (Ebert & Brune, 1997; Köhler et al., 2012; Schmitt-Wagner & Brune, 1999). In *O. disjunctus*, methanogenesis is concentrated in the AHG, where a thickened gut wall and elevated H₂ levels support hydrogenotrophic metabolism (Ceja-Navarro et al., 2019). Our detection of low-diversity archaeal assemblages in the anterior hindgut across passalid hosts are therefore consistent with a broader pattern in lignocellulose feeders. Bacteria, by contrast, integrate both host-controlled filtering and microhabitat structure, and thus carry most of the detectable phylogenetic signal.

### Effects of Host Phylogeny in the microbial communities of Passalidae

Patterns of parallel evolution between passalid phylogeny and microbiota composition may arise without strict vertical transmission when phylogenetically conserved host traits, such as the intestinal bauplan, physicochemical gradients, and immune filtering, act as ecological filters on a shared microbial pool, yielding microbial communities whose dissimilarities track host relatedness (Beza-Beza et al., 2024). Simulations and “common-garden” experiments across animals show that such host-trait filtering alone can generate phylosymbiotic patterns (Brooks et al., 2016). In insects, different distance metrics capture different aspects of this congruence: presence/absence-based unweighted UniFrac and Jaccard distances emphasize rare taxa and global topology, whereas abundance-weighted UniFrac and Bray-Curtis distances are more sensitive to dominant clades and localized subtree matches (Brooks et al., 2016; Tinker & Ottesen, 2020). Our passalid data conform to this expectation. Generally bacterial dendrograms show significant congruence with host trees at genus level under the more sensitive, abundance- and topology-aware metrics, with the exception of the matching cluster test under weighted UniFrac, whereas archaea do not (e.g. weighted UniFrac, Figs. 8,9). Our results are consistent with this framework but also show that phylosymbiosis in Passalidae is not uniform across domains or phylogenetic scales.

For bacteria, the Jaccard, Bray-Curtis, and unweighted UniFrac phylobiomes maintained relationships among closely related host species, while weighted UniFrac produced a different topology driven by the placement of *Passalus punctiger*. This indicates that bacterial phylosymbiosis is strongest for broad community membership and closely related host species but is more sensitive to abundance shifts in dominant taxa. For archaea, by contrast, the phylobiomes recovered deeper host tribal relationships more consistently but failed to recover similarity patterns among closely related host species. This contrast suggests that bacterial communities may retain finer-scale host-associated structure, whereas archaeal assemblages are constrained by broader host or gut-environmental features that operate at deeper phylogenetic levels.

This pattern is consistent with mixed-mode transmission – enabled by a combination of frass-based provisioning, gallery reuse, and substrate-mediated seeding – that stabilizes host-lineage-specific communities at deeper scales but permits ongoing horizontal recruitment and replacement at shallower ones, eroding strict concerted diversification. Termites exemplify this regime: multi-lineage host-symbiont co-speciation over tens of millions of years is accompanied by frequent host switching and mixed-mode transmission (Arora et al., 2023; Bourguignon et al., 2018). Passalids may therefore show phylosymbiosis without strict codiversification: host traits and social transmission can make microbiota composition and structure resemble host phylogeny even when individual microbial lineages are not vertically inherited with high fidelity.

Finally, our results clarify where passalids sit relative to other wood-feeding systems in terms of transmission and ecological assembly. Termites and their *Cryptocercus* ancestors rely on proctodeal trophallaxis to vertically transfer hindgut symbionts with high fidelity (Nalepa, 2015), a process quantified at ∼0.9 per protist lineage in *Reticulitermes grassei* (Michaud et al., 2020). Cerambycids, in contrast, often acquire dominant gut bacteria and fungi from oviposition sites and frass, with life-stage cores that track substrate rather than host phylogeny (Mason et al., 2019; Scully et al., 2013). Passalids appear to fall somewhere between these extremes. Adults process wood through a compartmentalized gut, then provision larvae with nitrogen- and energy-rich frass and triturated wood, which both nourish offspring and inoculate them with the adult microbiota (Ceja-Navarro et al., 2019; Mishima et al., 2016). Frass in galleries also harbors diverse defensive *Streptomyces* across broad spatial scales (Pessotti et al., 2021), pointing to shared colony-level reservoirs that, aside from serving a role in nest hygiene (Cole et al., 2021), are nonetheless open to environmental recruitment. Combined with the conserved AHG fibrolytic guild described by Schwarz et al. (2023) and the cross-species recurrence of key fiber-associated taxa in our dataset, these features support a scenario in which partner-fidelity feedback and conserved gut architecture generate deep-node bacterial phylosymbiosis in Passalidae, while more open, frass-based transmission and environmental admixture prevent the strict co-diversification characteristic of some termite clades (Brune & Dietrich, 2015).

## Conclusion

In sum, our study provides the first multi-species, phylogenetically-anchored view of gut microbiota structure in Passalidae. Passalid microbial communities show structuring and significant phylosymbiosis, indicating that host evolutionary history is a major determinant of gut microbiome assembly. Geographic locality contributes additional, but secondary, variation. These patterns, combined with detailed single-species work on fiber-associated microbiomes and compartmental physiology in *O. disjunctus*, establish Passalidae as a powerful comparative system for testing how social behavior, gut compartmentalization, and environmental assembly interact in the evolution of wood-feeding symbioses. Genome-resolved surveys across passalid lineages, explicit quantification of transmission routes, and compartment-resolved dual-domain profiling will be key next steps for resolving how much of the unclassified diversity in this group represents beetle-specific radiations analogous to termite “termite-group” clades, and for understanding how often evolution arrives at similar microbiota solutions to the problem of digesting wood.

## Acknowledgments

Ana Maestre-Guerra for specimens’ intestine removal and map elaboration. Doctors, Joshua Jones and Marisa Chase for their valuable comments on early drafts of the work. The participation of Larry Jiménez-Ferbans in this work was carried out within the framework of his sabbatical year, granted by the University of Magdalena.

## Data availability statement

Microbiome raw sequence data can be access in the National Center for Biotechnology information under BioProject PRJNA1509021. All processed data (phyloseq objects, host phylogenetic tree, randomly generated host phylogenetic trees, phylobiomes, and phylogentic vs phylobiome distance matrix) can be found at the Data Repository for the University of Minnesota (DOI:).

## References

Abdul Rahman, N., Parks, D. H., Willner, D. L., Engelbrektson, A. L., Goffredi, S. K., Warnecke, F., Scheffrahn, R. H., & Hugenholtz, P. (2015). A molecular survey of Australian and North American termite genera indicates that vertical inheritance is the primary force shaping termite gut microbiomes. Microbiome, 3(1), 5. 10.1186/s40168-015-0067-8

Arora, J., Buček, A., Hellemans, S., Beránková, T., Romero Arias, J., Fisher, B. L., Clitheroe, C., Brune, A., Kinjo, Y., Šobotník, J., & Bourguignon, T. (2023). Evidence of cospeciation between termites and their gut bacteria on a geological time scale. Proceedings of the Royal Society B: Biological Sciences, 290(2001), 20230619. 10.1098/rspb.2023.0619

Bauer, E., Lampert, N., Mikaelyan, A., Köhler, T., Maekawa, K., & Brune, A. (2015). Physicochemical conditions, metabolites and community structure of the bacterial microbiota in the gut of wood-feeding cockroaches (Blaberidae: Panesthiinae). FEMS Microbiology Ecology, 91(2), 1–14. 10.1093/femsec/fiu028

Beza-Beza, C. F., Jiménez-Ferbans, L., & McKenna, D. D. (2021). Historical biogeography of New World passalid beetles (Coleoptera, Passalidae) reveals Mesoamerican tropical forests as a center of origin and taxonomic diversification. Biogeography, 48, 2037–2052.

Beza-Beza, C. F., Wiegmann, B. M., Ware, J. A., Petersen, M., Gunter, N., Cole, M. E., Schwarz, M., Bertone, M. A., Young, D., & Mikaelyan, A. (2024). Chewing through challenges: Exploring the evolutionary pathways to wood-feeding in insects. BioEssays, 46(5), 2300241. 10.1002/bies.202300241

Boucher, S. (2006). Evolution et phylogénie des coléoptères Passalidae (Scarabaeoidea). Annales De La Société Entomologique De France, 41, 237–604. 10.1080/00379271.2005.10697444

Bourguignon, T., Lo, N., Dietrich, C., Šobotník, J., Sidek, S., Roisin, Y., Brune, A., & Evans, T. A. (2018). Rampant host switching shaped the termite gut microbiome. Current Biology, 28(4), 649–654.e2. 10.1016/j.cub.2018.01.035

Brooks, A. W., Kohl, K. D., Brucker, R. M., Van Opstal, E. J., & Bordenstein, S. R. (2016). Phylosymbiosis: Relationships and functional effects of microbial communities across host evolutionary history. PLOS Biology, 14(11), e2000225. 10.1371/journal.pbio.2000225

Brune, A., & Dietrich, C. (2015). The gut microbiota of termites: digesting the diversity in the light of ecology and evolution. Annual Review of Microbiology, 69(1), 145–166. 10.1146/annurev-micro-092412-155715

Callahan, B. J., McMurdie, P. J., Rosen, M. J., Han, A. W., Johnson, A. J. A., & Holmes, S. P. (2016). DADA2: High-resolution sample inference from Illumina amplicon data. Nature Methods, 13(7), 581–583. 10.1038/nmeth.3869

Callahan, B. J., Sankaran, K., Fukuyama, J. A., McMurdie, P. J., & Holmes, S. P. (2016). Bioconductor workflow for microbiome data analysis: From raw reads to community analyses. F1000Research, 5, 1492. 10.12688/f1000research.8986.2

Castillo, M. L., & Morón, M. A. (1992). Observaciones sobre la degradación de madera por algunas especies de Pasálidos (Coleoptera: Lamelicornia). Folia Entomológica Mexicana, 84, 35–44.

Castillo, M. L., & Reyes-Castillo, P. (2009). Passalidae, insects which live in decaying logs. In K. Del Claro, P. S. Oliveira, V. Rico-Gray, A. Ramirez, A. A. Almeida-Barbosa, A. Bonet, F. Rubio-Scarano, F. L. Consoli, F. J. Morales-Garzon, J. Naoki-Nakajima, J. A. Costello, M. Vinicius-Sampaio, M. Quesada, M. R. Morris, M. Palacios-Rios, N. Ramirez, O. Marcal Junior, R. H. Ferraz-Macedo, R. J. Marquis, … U. Luttege (Eds.), Torpical Biology and Conservation Managment: VII. International Commission of Tropical Biology and Natural Resources.

Ceja-Navarro, J. A., Karaoz, U., Bill, M., Hao, Z., White, R. A., Arellano, A., Ramanculova, L., Filley, T. R., Berry, T. D., Conrad, M. E., Blackwell, M., Nicora, C. D., Kim, Y.-M., Reardon, P. N., Lipton, M. S., Adkins, J. N., Pett-Ridge, J., & Brodie, E. L. (2019). Gut anatomical properties and microbial functional assembly promote lignocellulose deconstruction and colony subsistence of a wood-feeding beetle. Nature Microbiology, 4(5), 864–875. 10.1038/s41564-019-0384-y

Ceja-Navarro, J. A., Nguyen, N. H., Karaoz, U., Gross, S. R., Herman, D. J., Andersen, G. L., Bruns, T. D., Pett-Ridge, J., Blackwell, M., & Brodie, E. L. (2014). Compartmentalized microbial composition, oxygen gradients and nitrogen fixation in the gut of *Odontotaenius disjunctus*. The ISME Journal, 8(1), 6–18. 10.1038/ismej.2013.134

Chakraborty, A., Šobotník, J., Votýpková, K., Hradecký, J., Stiblik, P., Synek, J., Bourguignon, T., Baldrian, P., Engel, M. S., Novotný, V., Odriozola, I., & Větrovský, T. (2023). Impact of wood age on termite microbial assemblages. Applied and Environmental Microbiology, 89(5), e00361–23. 10.1128/aem.00361-23

Chao, A., & Chiu, C. (2016). Species richness: Estimation and comparison. In R. S. Kenett, N. T. Longford, W. W. Piegorsch, & F. Ruggeri (Eds.), Wiley StatsRef: Statistics Reference Online (1st ed., pp. 1–26). Wiley. 10.1002/9781118445112.stat03432.pub2

Cole, M. E., Ceja-Navarro, J. A., & Mikaelyan, A. (2021). The power of poop: Defecation behaviors and social hygiene in insects. PLOS Pathogens, 17(10), e1009964. 10.1371/journal.ppat.1009964

Dietrich, C., Köhler, T., & Brune, A. (2014). The cockroach origin of the termite gut microbiota: Patterns in bacterial community structure reflect major evolutionary events. Applied and Environmental Microbiology, 80(7), 2261–2269. 10.1128/AEM.04206-13

Ebert, A., & Brune, A. (1997). Hydrogen concentration profiles at the oxic-anoxic interface: A microsensor study of the hindgut of the wood-feeding lower termite *Reticulitermes flavipes* (Kollar). Applied and Environmental Microbiology, 63(10), 4039– 4046. 10.1128/aem.63.10.4039-4046.1997

Egert, M., Wagner, B., Lemke, T., Brune, A., & Friedrich, M. W. (2003). Microbial community structure in midgut and hindgut of the humus-feeding larva of *Pachnoda ephippiata* (Coleoptera: Scarabaeidae). Applied and Environmental Microbiology, 69(11), 6659–6668. 10.1128/AEM.69.11.6659-6668.2003

Fisher, R. A., Corbet, A. S., & Williams, C. B. (1943). The relation between the number of species and the number of individuals in a random sample of an animal population. The Journal of Animal Ecology, 12(1), 42. 10.2307/1411

Goluch, T., Bogdanowicz, D., & Giaro, K. (2020). VISUAL TREECMP: Comprehensive comparison of phylogenetic trees on the web. Methods in Ecology and Evolution, 11(4), 494–499. 10.1111/2041-210X.13358

Han, C.J., Cheng, C.-H., Yeh, T.-F., Pauchet, Y., & Shelomi, M. (2024). Coconut rhinoceros beetle digestive symbiosis with potential plant cell wall degrading microbes. Npj Biofilms and Microbiomes, 10(1), 34. 10.1038/s41522-024-00505-9

Hayward, A., Poulin, R., & Nakagawa, S. (2021). A broadscale analysis of host-symbiont cophylogeny reveals the drivers of phylogenetic congruence. Ecology Letters, 24(8), 1681–1696. 10.1111/ele.13757

Hernández-García, J. A., Gonzalez-Escobedo, R., Briones-Roblero, C. I., Cano-Ramírez, C., Rivera-Orduña, F. N., & Zúñiga, G. (2018a). Gut bacterial communities of *Dendroctonus valens* and *D. mexicanus* (Curculionidae: Scolytinae): A metagenomic analysis across different geographical locations in Mexico. International Journal of Molecular Sciences, 19(9), 2578. 10.3390/ijms19092578

Hervé, V., Liu, P., Dietrich, C., Sillam-Dussès, D., Stiblik, P., Šobotník, J., & Brune, A. (2020). Phylogenomic analysis of 589 metagenome-assembled genomes encompassing all major prokaryotic lineages from the gut of higher termites. PeerJ, 8, e8614. 10.7717/peerj.8614

Huang, X.-F., Chaparro, J. M., Reardon, K. F., Judd, T. M., & Vivanco, J. M. (2016). Supplementing blends of sugars, amino acids, and secondary metabolites to the diet of termites (*Reticulitermes flavipes*) drive distinct gut bacterial communities. Microbial Ecology, 72(3), 497–502. 10.1007/s00248-016-0792-y

Jácome-Hernández, A., Lamelas, A., Desgarennes, D., Huerta, C., Cruz-Rosales, M., & Favila, M. E. (2023). Influence of phylogenetic, environmental, and behavioral factors on the gut bacterial community structure of dung beetles (Scarabaeidae: Scarabaeinae) in a neotropical biosphere reserve. Frontiers in Microbiology, 14, 1224601. 10.3389/fmicb.2023.1224601

Jiménez-Ferbans, L., Maestre-Guerra, A., Villalba-Fuentes, E., Barros-Barrios, M. M., & Muñoz-Montero, J. (2023). Passalidae (Coleoptera, Scarabaeoidea) from the Caribbean coast of Colombia: Synopsis, key, and new species description. ZooKeys, 1179, 243–297. 10.3897/zookeys.1179.104037

Jiménez-Ferbans, L., Reyes-Castillo, P., & Schuster, J. C. (2018). Passalidae (Coleoptera: Scarabaeoidea) of the biogeographical province of Chocó and the west Andean region of Colombia, with the description of two new species. Neotropical Entomology, 47(5), 642–667. 10.1007/s13744-017-0584-1

Köhler, T., Dietrich, C., Scheffrahn, R. H., & Brune, A. (2012). High-resolution analysis of gut environment and bacterial microbiota reveals functional compartmentation of the gut in wood-feeding higher termites (*Nasutitermes* spp.). Applied and Environmental Microbiology, 78(13), 4691–4701. 10.1128/AEM.00683-12

Lemke, T., Stingl, U., Egert, M., Friedrich, M. W., & Brune, A. (2003). Physicochemical conditions and microbial activities in the highly alkaline gut of the humus-feeding larva of *Pachnoda ephippiata* (Coleoptera: Scarabaeidae). Applied and environmental microbiology, 69(11), 6650–6658. https://journals.asm.org/doi/10.1128/aem.69.11.6650-6658.2003

Lim, S. J., & Bordenstein, S. R. (2020). An introduction to phylosymbiosis. Proceedings of the Royal Society B: Biological Sciences, 287(1922), 20192900. 10.1098/rspb.2019.2900

Manjula, A., Pushpanathan, M., Sathyavathi, S., Gunasekaran, P., & Rajendhran, J. (2015). Comparative analysis of microbial diversity in termite gut and termite nest using ion sequencing. Current Microbiology. 10.1007/s00284-015-0947-y

Mason, C. J., Campbell, A. M., Scully, E. D., & Hoover, K. (2019). Bacterial and fungal midgut community dynamics and transfer between mother and brood in the Asian Longhorned Beetle (*Anoplophora glabripennis*), an invasive xylophage. Microbial Ecology, 77(1), 230–242. 10.1007/s00248-018-1205-1

McMurdie, P. J., & Holmes, S. (2013). phyloseq: An R package for reproducible interactive analysis and graphics of microbiome census data. PLoS ONE, 8(4), e61217. 10.1371/journal.pone.0061217

Michaud, C., Hervé, V., Dupont, S., Dubreuil, G., Bézier, A. M., Meunier, J., Brune, A., & Dedeine, F. (2020). Efficient but occasionally imperfect vertical transmission of gut mutualistic protists in a wood-feeding termite. Molecular Ecology, 29(2), 308–324. 10.1111/mec.15322

Mikaelyan, A., Dietrich, C., Köhler, T., Poulsen, M., Sillam-Dussès, D., & Brune, A. (2015). Diet is the primary determinant of bacterial community structure in the guts of higher termites. Molecular Ecology, 24(20), 5284–5295. 10.1111/mec.13376

Mikaelyan, A., Köhler, T., Lampert, N., Rohland, J., Boga, H., Meuser, K., & Brune, A. (2015). Classifying the bacterial gut microbiota of termites and cockroaches: A curated phylogenetic reference database (DictDb). Systematic and Applied Microbiology, 38(7), 472–482. 10.1016/j.syapm.2015.07.004

Mikaelyan, A., Meuser, K., & Brune, A. (2017). Microenvironmental heterogeneity of gut compartments drives bacterial community structure in wood- and humus-feeding higher termites. FEMS Microbiology Ecology, 93(1), fiw210. 10.1093/femsec/fiw210

Mikaelyan, A., Strassert, J. F. H., Tokuda, G., & Brune, A. (2014). The fibre-associated cellulolytic bacterial community in the hindgut of wood-feeding higher termites (*Nasutitermes* spp.). Environmental Microbiology, 16(9), 2711–2722. 10.1111/1462-2920.12425

Minh, B. Q., Schmidt, H. A., Chernomor, O., Schrempf, D., Woodhams, M. D., Von Haeseler, A., & Lanfear, R. (2020). IQ-TREE 2: New models and efficient methods for phylogenetic inference in the genomic era. Molecular Biology and Evolution, 37(5), 1530–1534. 10.1093/molbev/msaa015

Mishima, T., Wada, N., Iwata, R., Anzai, H., Hosoya, T., & Araya, K. (2016). Super-protective child-rearing by Japanese Bess Beetles, *Cylindrocaulus patalis*: Adults provide their larvae with chewed and predigested wood. Insects, 7(2), 18. 10.3390/insects7020018

Nalepa, C. A. (2015). Origin of termite eusociality: Trophallaxis integrates the social, nutritional, and microbial environments. Ecological Entomology, 40(4), 323–335. 10.1111/een.12197

Oksanen, J., Simpson, G., Blanchet, F., Kindt, R., Legendre, P., Minchin, P., O’Hara, R., Solymos, P., Stevens, M., Szoecs, E., Wagner, H., Barbour, M., Bedward, M., Bolker, B., Borcard, D., Borman, T., Carvalho, G., Chirico, M., De Caceres, M., … Weedon, J. (2025). vegan: An R package for community ecologists. https://vegandevs.github.io/vegan/

Pessotti, R. D. C., Hansen, B. L., Reaso, J. N., Ceja-Navarro, J. A., El-Hifnawi, L., Brodie, E. L., & Traxler, M. F. (2021). Multiple lineages of *Streptomyces* produce antimicrobials within passalid beetle galleries across eastern North America. eLife, 10, e65091. 10.7554/eLife.65091

Quast, C., Pruesse, E., Yilmaz, P., Gerken, J., Schweer, T., Yarza, P., Peplies, J., & Glöckner, F. O. (2012). The SILVA ribosomal RNA gene database project: Improved data processing and web-based tools. Nucleic Acids Research, 41(D1), D590–D596. 10.1093/nar/gks1219

Schmitt-Wagner, D., & Brune, A. (1999). Hydrogen profiles and localization of methanogenic activities in the highly compartmentalized hindgut of soil-feeding higher termites (*Cubitermes* spp.). Applied and Environmental Microbiology, 65(10), 4490– 4496. 10.1128/AEM.65.10.4490-4496.1999

Schwarz, M., Beza-Beza, C. F., & Mikaelyan, A. (2023). Wood fibers are a crucial microhabitat for cellulose- and xylan-degrading bacteria in the hindgut of the wood-feeding beetle *Odontotaenius disjunctus*. Frontiers in Microbiology, 14, 1173696. 10.3389/fmicb.2023.1173696

Scully, E. D., Geib, S. M., Hoover, K., Tien, M., Tringe, S. G., Barry, K. W., Glavina Del Rio, T., Chovatia, M., Herr, J. R., & Carlson, J. E. (2013). Metagenomic profiling reveals lignocellulose degrading system in a microbial community associated with a wood-feeding beetle. PLoS ONE, 8(9), e73827. 10.1371/journal.pone.0073827

Su, L., Yang, L., Huang, S., Su, X., Li, Y., Wang, F., Wang, E., Kang, N., Xu, J., & Song, A. (2016). Comparative gut microbiomes of four species representing the higher and the lower termites. Journal of Insect Science, 16(1), 97. 10.1093/jisesa/iew081

Tinker, K. A., & Ottesen, E. A. (2020). Phylosymbiosis across deeply diverging lineages of omnivorous cockroaches (Order Blattodea). Applied and Environmental Microbiology, 86(7), e02513–19. 10.1128/AEM.02513-19

Tokuda, G., Mikaelyan, A., Fukui, C., Matsuura, Y., Watanabe, H., Fujishima, M., & Brune, A. (2018). Fiber-associated spirochetes are major agents of hemicellulose degradation in the hindgut of wood-feeding higher termites. Proceedings of the National Academy of Sciences, 115(51). 10.1073/pnas.1810550115

Wang, Y., & Rozen, D. E. (2017). Gut microbiota colonization and transmission in the burying beetle *Nicrophorus vespilloides* throughout development. Applied and Environmental Microbiology, 83(9), e03250–16. 10.1128/AEM.03250-16

