## Supplementary Files for "Termite-Like Phylogenetic Conservation of Gut Microbiota in Passalid Beetles Suggests Evolutionary Convergence in Decaying Wood Degradation"

Supplementary Material list.

Tables

Supplementary table S1. Detailed specimen information.

| Species | Colony ID | Geographical coordinates | Altitude (msnm) | Total Colony individuals (# sequenced individuals) | Microbial Sequence ID |
|---|---|---|---|---|---|
| **Sierra Nevada de Santa Marta** | | | | | |
| *Passalus* (*Passalus*) *coniferus* | 1 | 11°04’02” N 74°02’41” W | 1662 | 1(1) | SM10 |
| *Passalus* (*Passalus*) *punctiger* | 1 | 11°04’02” N 74°02’41” W | 1662 | 2(2) | SM8 |
| *Passalus* (*Passalus*) *punctiger* | 3 | 11°04’02” N 74°02’41” W | 1662 | 2(2) | SM9 |
| *Passalus* (*Pertinax*) *unimagdalenae* | 1 | 11°04’02” N 74°02’41” W | 1662 | 6(4) | SM11, SM12 |
| *Passalus* (*Pertinax*) *unimagdalenae* | 2 | 11°04’02” N 74°02’41” W | 1662 | 4(2) | SM13 |
| *Popilius marginatus* | 1 | 11°04’02” N 74°02’41” W | 1662 | 17(4) | SM2, SM3 |
| *Popilius marginatus* | 2 | 11°04’02” N 74°02’41” W | 1662 | 8(4) | SM4, SM5 |
| *Popilius marginatus* | 3 | 11°04’02” N 74°02’41” W | 1662 | 8(4) | SM6, SM7 |
| *Veturius impressus* | 1 | 11°03’56” N 74°03’05” W | 1684 | 5(4) | SM14, SM15 |
| *Veturius impressus* | 2 | 11°03’56” N 74°03’05” W | 1684 | 2(2) | SM16 |
| *Veturius impressus* | 4 | 11°04’02” N 74°02’41” W | 1662 | 2(2) | SM1 |
| *Veturius impressus* | 5 | 11°03’56” N 74°03’05” W | 1684 | 4(4) | SM17, SM18 |
| **Serranía del Perijá** | | | | | |
| *Rodocanthopus maillei* | 1 | 10°50'15.5"N 72°40'04.8"W | 1785 | 9(4) | P7, P8 |
| *Rodocanthopus maillei* | 2 | 10°50'15.8"N 72°40'12.4"W | 1745 | 11(4) | P1, P2 |
| *Rodocanthopus maillei* | 3 | 10°50'12.8" N 72°40'01.6"W | 1792 | 4(2) | P13 |

| | | | | | |
|---|---|---|---|---|---|
| *Popilius gibbosus* | 1 | 10°50'16.1"N, 72°40'10.5"W | 1777 | 11(4) | P5, P6 |
| *Popilius gibbosus* | 2 | 10°50'12.8" N 72°40'01.6"W | 1792 | 3(2) | P12 |
| *Veturius standfussi* | 1 | 10°50'15.8"N 72°40'12.4"W | 1745 | 5(4) | P3, P4 |
| *Veturius standfussi* | 2 | 10°50'12.8" N 72°40'01.6"W | 1792 | 5(4) | P10, P11 |

Supplementary table S2. Read count and DADA2 processing for Bacteria.

| Sample | input | filtered | denoisedF | denoisedR | merged | nonchim |
|---|---|---|---|---|---|---|
| **P10a** | 205583 | 183626 | 182377 | 181769 | 173658 | 140786 |
| **P11a** | 206944 | 170709 | 169844 | 169206 | 159967 | 123784 |
| **P12a** | 204266 | 175141 | 173909 | 173116 | 164339 | 139471 |
| **P13a** | 202332 | 173451 | 172665 | 172467 | 168399 | 147603 |
| **P1a** | 213066 | 190763 | 189867 | 189672 | 184789 | 164400 |
| **P2a** | 204826 | 179554 | 178386 | 177616 | 170546 | 141144 |
| **P3a** | 203620 | 169119 | 168091 | 167812 | 161767 | 122223 |
| **P4a** | 212325 | 188056 | 186865 | 186164 | 179365 | 124854 |
| **P5a** | 219766 | 184018 | 183834 | 183749 | 182577 | 85414 |
| **P6a** | 206172 | 174043 | 173192 | 172360 | 165611 | 128440 |
| **P7a** | 204151 | 170038 | 169054 | 168902 | 164248 | 140227 |
| **P8a** | 204371 | 182228 | 181328 | 181341 | 176911 | 148337 |
| **SM10a** | 217214 | 184606 | 183266 | 183515 | 178238 | 130036 |
| **SM11a** | 205529 | 182022 | 180653 | 180335 | 172701 | 149024 |
| **SM12a** | 202940 | 181488 | 180176 | 179851 | 173072 | 137111 |
| **SM13a** | 205941 | 181741 | 180547 | 180117 | 173067 | 152153 |
| **SM14a** | 204177 | 185405 | 184085 | 183446 | 175334 | 145643 |
| **SM15a** | 169635 | 155584 | 154737 | 154491 | 149613 | 113074 |
| **SM16a** | 204489 | 182879 | 182042 | 181868 | 175830 | 160808 |
| **SM17a** | 203834 | 185229 | 183829 | 183647 | 176780 | 133509 |
| **SM18a** | 173966 | 158674 | 157658 | 157432 | 152449 | 124464 |
| **SM1a** | 205308 | 183531 | 182668 | 182699 | 178109 | 163175 |

| | | | | | | |
|---|---|---|---|---|---|---|
| **SM2a** | 202706 | 183122 | 182511 | 182105 | 177973 | 158746 |
| **SM3a** | 218583 | 197237 | 196129 | 195685 | 188885 | 164149 |
| **SM4a** | 203498 | 178348 | 177432 | 176722 | 170042 | 147940 |
| **SM5a** | 219630 | 196103 | 194113 | 193499 | 181659 | 150343 |
| **SM6a** | 205245 | 186512 | 185929 | 185778 | 181875 | 167490 |
| **SM7a** | 202169 | 181276 | 180582 | 180326 | 175920 | 160819 |
| **SM8a** | 209235 | 187949 | 187347 | 187241 | 181379 | 145341 |
| **SM9a** | 203091 | 182999 | 181162 | 180873 | 171706 | 137795 |

Supplementary table S3. Read count and DADA2 processing for Archaea.

| Sample | input | filtered | denoisedF | denoisedR | merged | nonchim |
|---|---|---|---|---|---|---|
| **P10b** | 201121 | 186972 | 183028 | 183737 | 145321 | 93266 |
| **P11b** | 122659 | 114384 | 112136 | 112180 | 91476 | 64664 |
| **P13b** | 202991 | 188523 | 184617 | 185366 | 156361 | 114406 |
| **P2b** | 202645 | 186512 | 180438 | 181782 | 136708 | 95659 |
| **P3b** | 108841 | 101405 | 100162 | 99949 | 85627 | 64291 |
| **P4b** | 206616 | 191214 | 187652 | 188379 | 157840 | 125565 |
| **P5b** | 209602 | 193614 | 188401 | 189755 | 147353 | 98914 |
| **P6b** | 152334 | 141150 | 136126 | 136915 | 99903 | 60573 |
| **P7b** | 169511 | 157497 | 156209 | 155490 | 141439 | 118943 |
| **P8b** | 206700 | 191901 | 189064 | 189828 | 168238 | 135700 |
| **SM10b** | 202245 | 187834 | 184434 | 185047 | 155721 | 119664 |
| **SM11b** | 203933 | 187788 | 182712 | 183760 | 140398 | 95057 |
| **SM12b** | 202338 | 186593 | 180825 | 182123 | 137220 | 92418 |
| **SM13b** | 206435 | 190766 | 186447 | 187317 | 147873 | 104987 |
| **SM14b** | 218240 | 201917 | 199573 | 200252 | 180536 | 147458 |
| **SM15b** | 204250 | 189142 | 185773 | 186514 | 154089 | 115554 |
| **SM16b** | 205129 | 189336 | 185740 | 186772 | 156714 | 111184 |
| **SM17b** | 204790 | 190106 | 185764 | 186730 | 153765 | 110967 |
| **SM18b** | 204442 | 189643 | 183985 | 185066 | 142053 | 98158 |
| **SM1b** | 205626 | 189329 | 183371 | 184089 | 130402 | 74214 |
| **SM2b** | 216030 | 200163 | 196265 | 197070 | 164012 | 102529 |
| **SM3b** | 202083 | 187200 | 182027 | 183177 | 139729 | 82155 |

| | | | | | | |
|---|---|---|---|---|---|---|
| **SM4b** | 205598 | 190025 | 185624 | 186601 | 147580 | 96395 |
| **SM5b** | 204097 | 189484 | 183734 | 184934 | 139127 | 83206 |
| **SM6b** | 203014 | 187761 | 182393 | 183275 | 139140 | 77904 |
| **SM7b** | 113193 | 104854 | 101733 | 102341 | 75861 | 44464 |
| **SM8b** | 141178 | 131058 | 128155 | 128726 | 106144 | 81230 |
| **SM9b** | 206236 | 190189 | 186631 | 187177 | 155984 | 111280 |

Supplementary table S4. Unclassified ASVs at different taxonomic levels, number in parentheses represent the percentage of total reads

| Rank | Bacteria ASVs (% reads) | Archaea ASVs (% reads) |
|---|---|---|
| **Class** | 128 (0.7 %) | 1 (< 0.01 %) |
| **Order** | 702 (3.9 %) | 4 (< 0.01 %) |
| **Family** | 3059 (10.4 %) | 18 (0.3 %) |
| **Genus** | 9628 (50.5 %) | 112 (2.71 %) |

Supplementary table S5. P values of Beta dispersion test at the level of tribe, genus, species and locality, under Jaccard, Bray-Curtis, and unweighted Unifrac

| | | Jaccard | Bray-Curtis | Unifrac | Weighted Unifrac |
|---|---|---|---|---|---|
| **Bacteria** | **Tribe** | 0.0001 | 0.0006 | 0.0000 | 0.135 |
| | **Genus** | 0.2442 | 0.4062 | 0.2187 | 0.7123 |
| | **Species** | 0.0000 | 0.0000 | 0.0000 | 0.1086 |
| | **Locality** | 0.4756 | 0.2532 | 0.6888 | 0.162 |
| **Arachaea** | **Tribe** | 0.049 | 0.214 | 0.020 | 0.000 |
| | **Genus** | 0.006 | 0.066 | 0.159 | 0.148 |
| | **Species** | 0.000 | 0.071 | 0.001 | 0.503 |
| | **Locality** | 0.004 | 0.132 | 0.047 | 0.145 |

Supplementary. Table S6. Summary statistics of normalized Robinson-Foulds (RF), and Matching Cluster (MC) distances between host phylogeny and 1000 randomly generated phylogenetic trees

| **Metric** | **Mean** | **sd** |
|---|---|---|
| **RF** | 0.896 | 0.096 |
| **MC** | 0.817 | 0.118 |

Figures

Figure S1. Heat map of relative abundance of the Top 20 Archaeal ASVs.

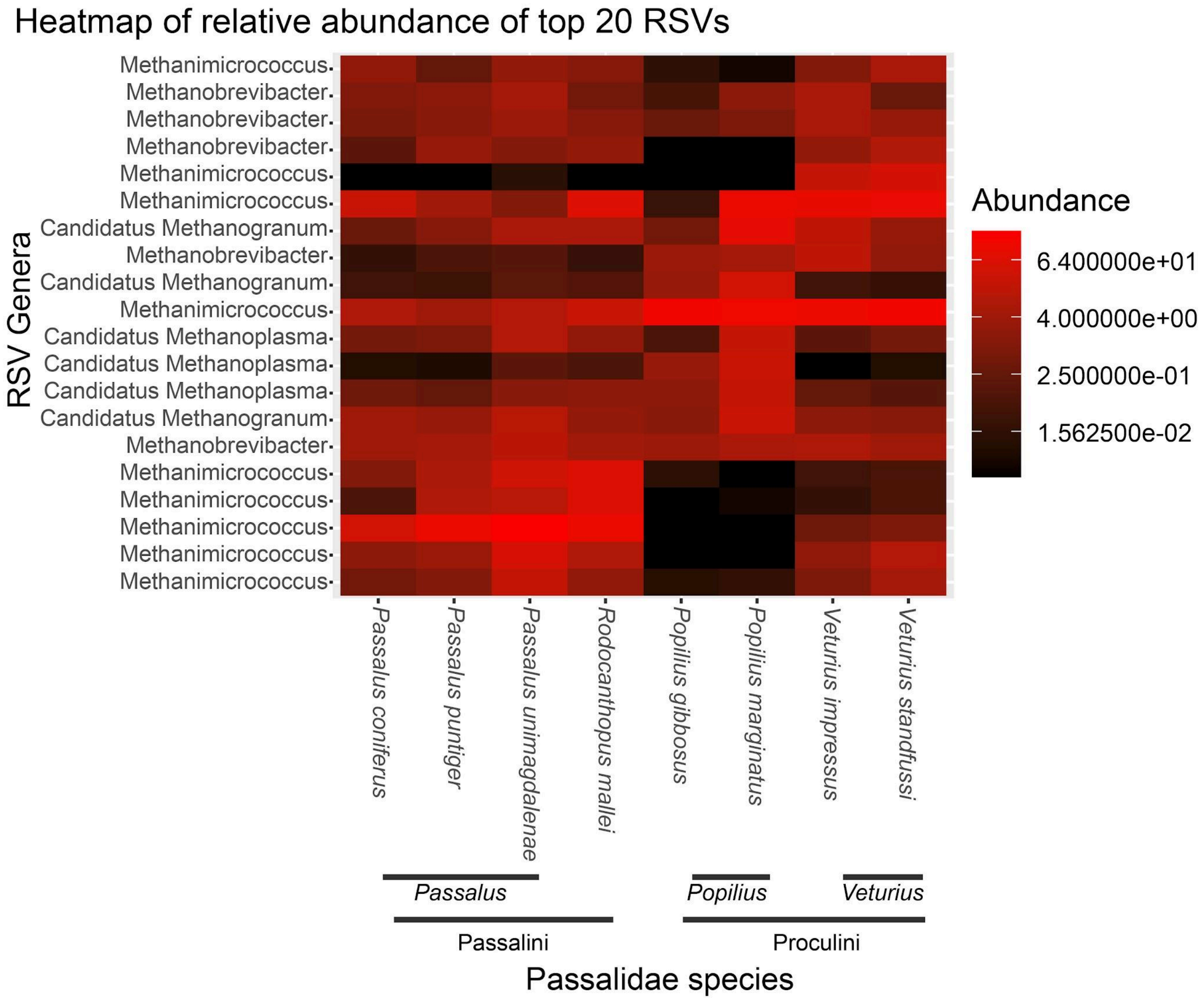
